# Programmable viscoelastic hydrogels uncover mechano-sensing timescales and direct cell polarity

**DOI:** 10.1101/2025.02.09.637331

**Authors:** Syuan-Ku Hsiao, Markus Mukenhirn, Carsten Werner, Alf Honigmann, Elisha Krieg

## Abstract

Mechanical interactions between cells and their three-dimensional environment govern fundamental processes in tissue development and disease. Yet, how cells interrogate and respond to mechanical cues remains incompletely understood. Reproducing the mechanical properties of natural tissues *in vitro* is a key challenge, as these tissues exhibit complex viscoelastic behaviors and undergo continuous remodeling throughout an organism’s development. Here, we leverage principles of dynamic DNA nanotechnology to introduce programmable mechanical cues into a synthetic hydrogel matrix that guides and interrogates the development of embedded cells. By systematically modulating matrix stress relaxation and stiffness, we uncover *two distinct timescales* of mechano-sensitive processes controlling apical–basal polarity in Madin–Darby Canine Kidney (MDCK) cells: the fast timescale (1–3 min) is linked to rapid integrin-mediated signaling, while the slower process (3–9 h) is associated with cytoskeletal reinforcements. Our analysis highlights that the commonly reported matrix “stiffness” value, often measured as the storage modulus at ∼1 Hz, has limited physiological relevance. These findings prompted us to develop a novel *switchable DNA crosslinker* that enables dynamic changes in viscoelasticity during ongoing cell culture. This controlled matrix reconfiguration induces reversible cell polarity inversions and guides the morphogenesis of complex multicellular structures. The material platform therefore offers unprecedented control over the mechanical microenvironment, opening new avenues for advanced applications in biophysics, tissue engineering, and disease modeling.

## Introduction

The development of biological tissues is controlled by mechano-chemical interactions between cells and the *extracellular matrix* (*ECM*)^1–3^. The transduction of mechanical signals across the cell membrane through *integrin receptors* is crucial for sensing and cell decision-making in all multicellular organisms^4,5^. Cell polarization is one of the fundamental processes that largely depend on mechano-signaling^6–8^. In epithelial monolayers, polarization establishes distinct *apical* and *basolateral* membrane domains^9^. Mechano-chemical stimulation at the basal interface provides a key signal for the alignment of the apical interface “away” from the basal side. Ion transport across the apical membrane creates osmotic gradients that drive the expansion of a lumen, the interior space of the hollow multicellular structure^10^. The apical side facilitates critical functions such as nutrient uptake, protein reabsorption, and interactions with commensal bacteria or pathogens^11,12^. The malfunction of cellular mechano-signaling is therefore associated with a wide range of human diseases, including rare genetic disorders and various types of cancer^2,4,5,13–15^.

Seminal research has yielded important insights into mechano-chemical ECM signaling in cells grown on two-dimensional surfaces^16^ and in more realistic three-dimensional (3D) environments^12,17^. However, the specific mechanical properties of the ECM that cells sense—and how they do so—remain open questions. A growing body of research indicates that cells sense more than just the stiffness of their environment^18–23^. In particular, time-dependent matrix stress relaxation, plasticity, and non-linear deformation characteristics have been shown to play important roles in cellular mechano-signaling^17,24^. To systematically study these dynamic properties in a realistic *in vitro* environment, novel 3D cell culture matrices with customizable mechanical and biochemical properties are needed^24–29^.

Developing more realistic 3D matrices with precisely adjustable mechanical properties is not a trivial task. To control matrix stress relaxation and plasticity, one must engineer the timescale of conformational rearrangements and crosslinker breakage in response to mechanical deformation. Notably, Chaudhuri and colleagues reported hydrogels based on alginate, whose molecular weight and functionalization changed the material’s stress relaxation^21–23^. Others reported the use of dynamic covalent bonds and supramolecular motifs to modulate crosslinker stability^18–20^. However, there remain significant challenges for precisely adjusting a material’s properties without introducing confounding changes to its chemical functionalization or the medium’s composition. This engineering problem is further complicated by the dynamic nature of biological tissues: unlike synthetic materials, biological matter is continuously remodeled. Changes in mechanical properties are part of the organism’s healthy development, but they can also be associated with dysfunctions and diseases. There is therefore a large need for a next generation of materials with *programmable* and *reconfigurable* mechanical characteristics.

DNA nanotechnology^30^ offers an intriguing approach to this challenge. DNA molecules can be designed to self-assemble into distinct nanoscopic objects with custom size, shape, and programmable dynamic transformations. DNA nanosystems have been primarily applied in solution or on surfaces; however, they can also be used to engineer soft hydrogels that provide a high level of control from nanoscale structure to macroscale behavior^31–36^.

In this study, we developed an evolved variant of a DNA-crosslinked hydrogel matrix (*DyNAtrix*^36^) to explore how changes in matrix viscoelasticity influence apical–basal polarization in Madin– Darby Canine Kidney (MDCK) cysts, a popular organotypic model for epithelial cell polarity. We introduced a new correlation-based data analysis that links systematic changes in matrix stress-relaxation to the timescales of mechano-signaling events. This approach enabled us to identify two distinct timescales underlying the molecular mechanisms of cell polarization and lumen formation. Building on the striking discovery that tissue polarization can be regulated by tuning stress relaxation, we went one step further and developed a conceptually new generation of switchable crosslinker modules. For the first time, these modules allow the reversible modulation of matrix stress relaxation during an ongoing cell culture. The switchable stress relaxation in this advanced *DyNAtrix* variant provides a novel mechanism to actively guide cell-fate decisions through changes in integrin-mediated mechanical cell–matrix tension.

## Results

### Control over cell–matrix adhesion, stiffness, and stress relaxation

A set of *DyNAtrix* gels with different properties was produced through self-assembly of a poly(acylamide-co-acrylic acid)^37,38^ backbone with DNA-based crosslinker modules (Figure 1). The main backbone derivative (**P**^**+RGD**^) was furnished with peptide side chains containing the *RGD* (*Arg-Gly-Asp*) cell adhesion motif. A second backbone derivative (**P**) was synthesized under identical conditions but without RGD groups. **P** was used to assemble control gels that lack cell adhesion sites. The backbone was crosslinked through previously described combinatorial dual-splint^36^ DNA sequences that self-assemble with *anchor* strands on the polymer backbone (Figure 1a). One splint strand (*crosslinker 1*) was annealed to a complementary blocking strand (*blocker 1*). The second splint strand (*crosslinker 2*) was annealed to its respective blocking strand (*blocker 2*) and to the polymer backbone (see Methods section). These two precursor solutions were mixed at 4°C without premature crosslinking, as the blocking strands inhibit hybridization between the crosslinkers^36^. Subsequent heating to 37°C dissociated the blocking strands and thus triggered gelation without the need for chemical crosslinkers or photoactivation. This heat-activated gelation procedure allows gentle encapsulation of cells, mimicking the gelation behavior of widely used basement membrane matrices such as *Matrigel*.

**Figure 1.**
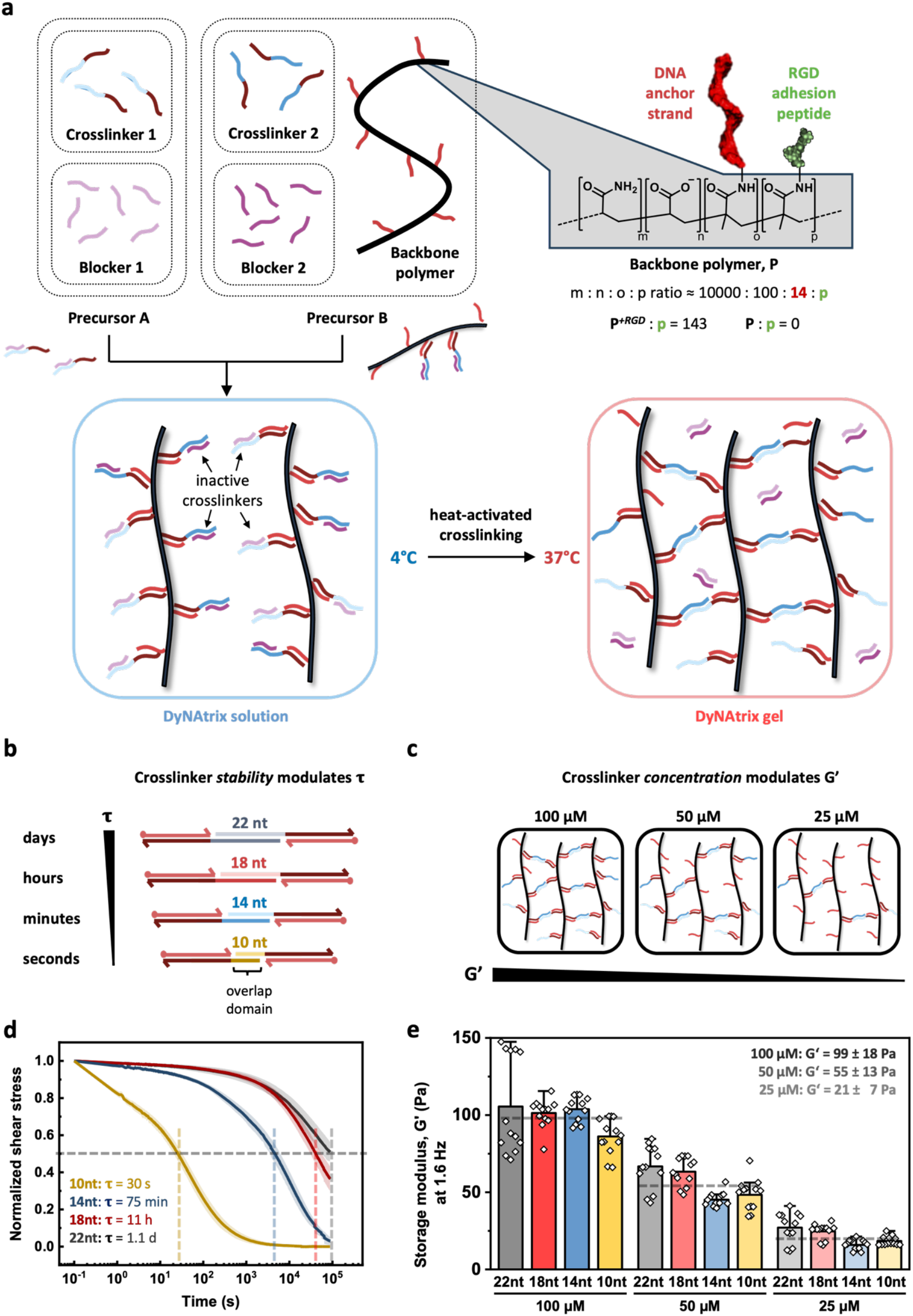
*DyNAtrix* stiffness and stress relaxation are adjustable via the crosslinker’s nanomechanical *stability* and its *concentration*. **a**, Scheme showing the structure of the *DyNAtrix* backbone polymer, crosslinker modules, and the pathway for matrix assembly. For clarity, RGD peptides are omitted from the crosslinking scheme. Black: polymer backbone; red: anchor strands; blue: overlap domains, purple: blocking strands. **b**, The set of four stress-relaxation crosslinkers (SRCs) that were used to generate a wide range of stress-relaxation characteristics. The SRCs are held together by *overlap domains* with different nanomechanical stabilities. The overlap domains are designed to break reversibly under mechanical tension, thus providing a mechanism for attenuating mechanical forces within the gel’s network. Overlap domains with descending stability are drawn in black, red, blue, and yellow. **c**, *DyNAtrix*’s shear elastic modulus (G’) is adjusted via the crosslinker concentration (100 µM, 50 µM or 25 µM). **d**, Normalized stress-relaxation curves of *DyNAtrix* crosslinked with the four different SRCs (at strain γ = 15%). The gel’s stress-relaxation time (**τ**) is defined as the time needed for the shear stress to decay to half of its initial value. The shaded area indicates the standard deviation of the stress-relaxation profiles obtained from three different samples of identical crosslinker stability but different crosslinker concentrations (see Supplementary Figure S3). **e**, Comparison of the storage moduli of *DyNAtrix* crosslinked with different SRCs and crosslink concentrations. Average values and standard deviations are based on at least 12 measurement replicates.

Four pairs of crosslinking strands were designed, varying only by the number of nucleotides (nt) in their *overlap* domains (Figure 1b). The mechanical force required to break a short DNA duplex increases with increasing duplex length^39^. As the overlap domain represents the weakest link in the supramolecular network, its predictable nanomechanical stability defines the gel’s stress-relaxation behavior^36^. The set of these so-called *stress-relaxation crosslinkers* (*SRC*) with 10-, 14-, 18-, or 22-nt overlap domains are denoted **SRC-10, SRC-14, SRC-18**, and **SRC-22**, respectively. Step-strain experiments demonstrated that the four alternative SRCs allowed precise adjustment of the material’s stress-relaxation time (τ) over four orders of magnitude, ranging from the time scale of seconds to the time scale of days (Figure 1d).

While the relaxation time was controlled through crosslinker *stability*, changes in the crosslinker *concentration* provided control over the material’s storage modulus (G’) (Figure 1c). The average measured G’ values at 1.6 Hz (i.e., the high-frequency plateau moduli) were 99±18 Pa, 55±13 Pa, and 21±7 Pa for gels that were prepared with 100 µM, 50 µM, and 25 µM crosslinker strands, respectively (Figure 1e). Expectedly, the measured plateau moduli were relatively independent of the changes in mechanical stabilities of the SRCs. Conversely, the stress-relaxation times were relatively insensitive to variations in crosslinker concentration (Figure 1d, Supplementary Figure S3). Thus, *DyNAtrix* allowed straightforward tuning of two important mechanical parameters, G’ and τ, without the need for synthetic modifications or changes in the medium’s composition.

### Cell polarization depends on cell adhesion, matrix stiffness and relaxation

To explore how mechanical anchoring and matrix viscoelasticity affect cell polarization, we first embedded MDCK cells in *DyNAtrix* with either 100 µM or 50 µM **SRC-14** that was assembled with either **P**^**+RGD**^ or **P** (Figure 2a–c). The cells were engineered to express fluorescently labeled *podocalyxin* and *E-cadherin* for easy identification of cell polarity and adherens junctions, respectively. To quantify cyst polarization, we group the observed morphologies into three classes: apical-in with a central lumen (*type I*), apical-out without lumen (*type II*), and apical-out with lumen (*type III*) (Figure 2a). Nearly all cysts grown in *DyNAtrix* with **P** backbone exhibited non-physiological apical-out morphologies, irrespective of the stiffness or stress-relaxation behavior of the matrix (Figure 2b). Thus, the presence of integrin-matrix adhesion sites was essential for the cells to develop correct apical-in polarity. This finding confirms that apical–basal polarization is controlled by specific integrin-mediated mechanical cues^8,12,40–43^ and further demonstrates that in the absence of integrin binding, cell polarization is insensitive to varying non-specific mechanical parameters.

**Figure 2.**
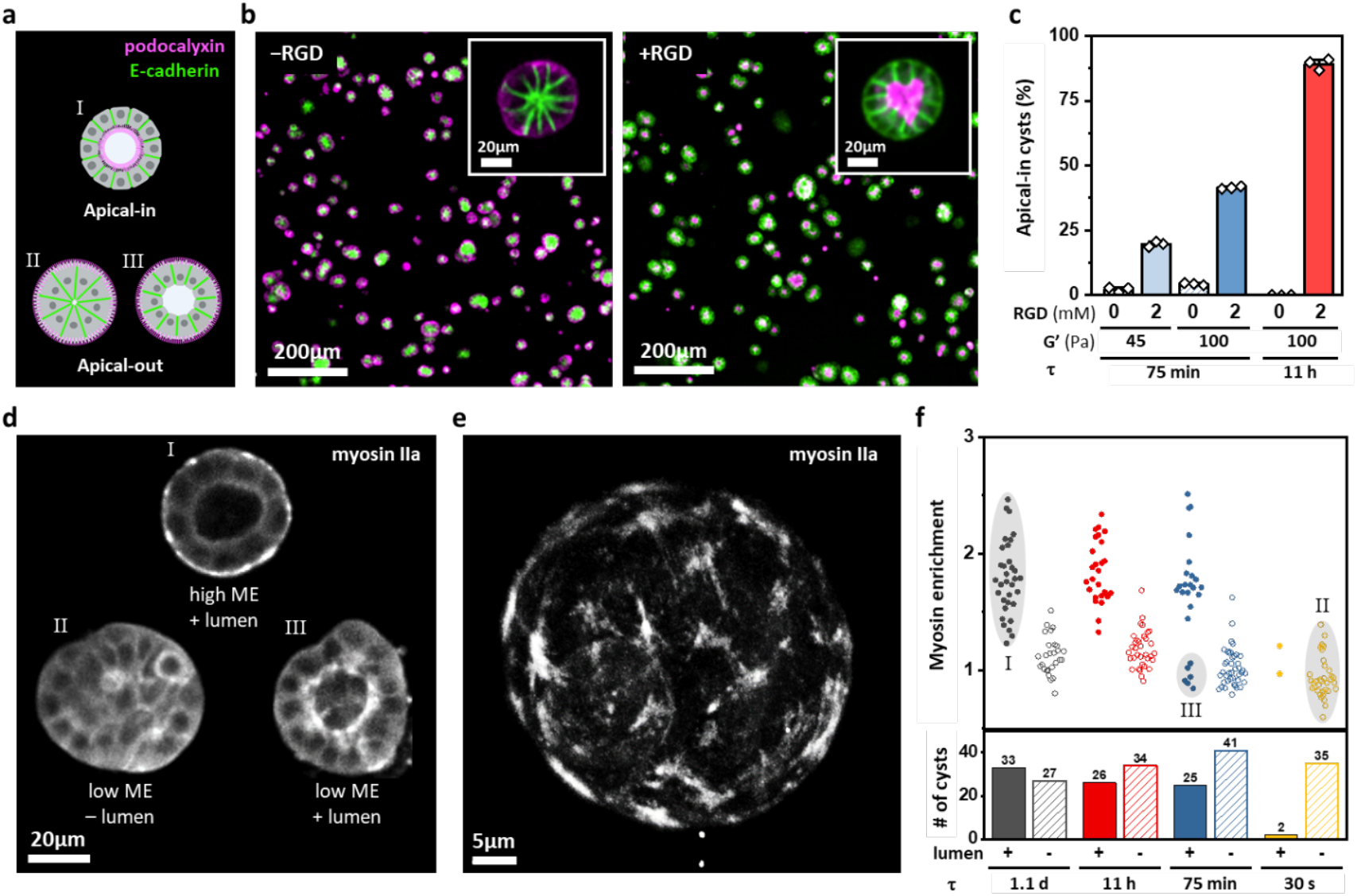
Mechanical cell-matrix interactions determine lumen formation and polarization, yielding three different types of MDCK cysts. **a**, Scheme illustrating the polarity of apical-in (top) versus apical-out (bottom) MDCK cysts. **b**, Confocal images of MDCK cysts (day 6) cultured in *DyNAtrix* (adjusted to G’ ∼100 Pa, τ ∼11 h) with versus without RGD. **c**, The fraction of apical-in cysts (on day 6) cultured in *DyNAtrix* with different concentrations of RGD, different stiffnesses, and different stress-relaxation times. **d**, Confocal images (cross-section) of the three different types of cysts highlighted in (d) (see also Supplementary Figure S5). Top: type I cyst with lumen and elevated ME; bottom left: type II cyst with low ME, lacking a well-defined lumen; bottom right: type III cyst with low ME value but exhibiting a lumen. **e**, Representative confocal image (maximum projection) of a cyst (day 6) cultured in stiff and slow-relaxing *DyNAtrix* (G’ ∼100 Pa, τ ∼1.1 d). The clusters of myosin indicate focal adhesions. **f**, Myosin enrichment (ME) and lumen formation analysis of MDCK cysts (day 6) cultured in *DyNAtrix* (G’ ∼100 Pa) with the four different SRCs. Longer stress-relaxation times result in higher ME and more pronounced lumen formation. Three different groups of cysts exhibiting distinct morphologies are labeled (I, II, III).

In contrast, when the **P**^**+RGD**^ backbone was used, cell polarity strongly depended on both stiffness and stress relaxation. Only ∼20% apical-in cysts were observed in soft and fast-relaxing *DyNAtrix* (50 µM **SRC-14**, G’ ∼45 Pa, τ ∼75 min), as opposed to 40% in stiff and fast-relaxing *DyNAtrix* (100 µM **SRC-14**, G’ ∼100 Pa, τ ∼75 min). Further drastic changes in cyst polarity were observed when the material’s stress-relaxation behavior was altered without changing the stiffness: 85% of cysts exhibited apical-in morphology when grown in stiff and slow-relaxing *DyNAtrix* (100 µM **SRC-18**, G’∼100 Pa, τ ∼11 h). These initial observations prompted us to explore the effects of matrix viscoelasticity on cyst morphology over a wide range of systematically varied parameters.

To further examine the role of stress relaxation time on cyst polarization, we cultured MDCK cysts in *DyNAtrix*, utilizing a wide range of stress-relaxation times at 100 µM SRC concentration (G’ ∼100 Pa). In addition to polarization and lumen formation, we quantified the enrichment of endogenously labeled myosin IIa at the cyst–matrix interface, as the indicator for mechanical engagement of focal adhesions with the matrix^44^ (Figure 2d-f, Supplementary Figures S4, S5). In **slow**-relaxing *DyNAtrix* cultures (τ ∼11 h or 1.1 d), the fraction of apical-in polarized cysts with a central lumen (*type I*) showed a pronounced myosin enrichment at the cyst-matrix interface (Figure 2d and f). Here, myosin formed clusters at the basal interface that are typical of mechanically loaded focal adhesions (Figure 2e). These cultures also contained a small fraction of lumen-less cysts with apical-out polarization (*type II*), which showed no significant myosin enrichment. In **fast**-relaxing *DyNAtrix* (τ ∼30 s), type II cysts represented the vast majority of observed morphologies, indicating that the mechanical tension at the cell–matrix interface must be sustained for a critical amount of time to enable correct polarization and lumen formation.

Intriguingly, in *DyNAtrix* cultures with **intermediate** stress-relaxation (τ ∼75 min), we observed a sub-population of exotic *type III* cysts containing a central lumen but lacking basal myosin enrichment (Figure 2d, Supplementary Figure S5). It remains to be shown whether type III cysts are a distinct phenotype or rather represent a transition state between type I and II cysts. However, the observation that lumen formation and cell polarization may be triggered independently from each other supports the hypothesis that the underlying mechano-signaling mechanisms are not identical for both processes. Overall, the effect of the *time scale* of sustained mechanical tension adds another facet to the previously described biophysical and biochemical conditions affecting epithelial tissue development^45^.

### Viscoelasticity tuning exposes critical time scales of mechano-signaling

To better understand the effect of mechanical cell-matrix interactions on polarization, we embedded MDCK cells in *DyNAtrix* variants exhibiting all 12 possible combinations of stiffness and stress-relaxation. The cysts were cultured for up to 11 days (Figure 3a,b; Supplementary Figures S6–8). The experiments confirmed the trend that increasing stiffness and slower stress relaxation resulted in increasing fractions of apical-in cysts. Conversely, there is a trade-off between the two parameters, G’ and τ: a matrix with high stiffness combined with fast stress relaxation can yield similar fractions of apical-in cysts as a matrix with low stiffness and slow stress relaxation (e.g., 100 µM **SRC-14** yields similar apical-in cyst polarization numbers as 50 µM **SRC-22**).

**Figure 3.**
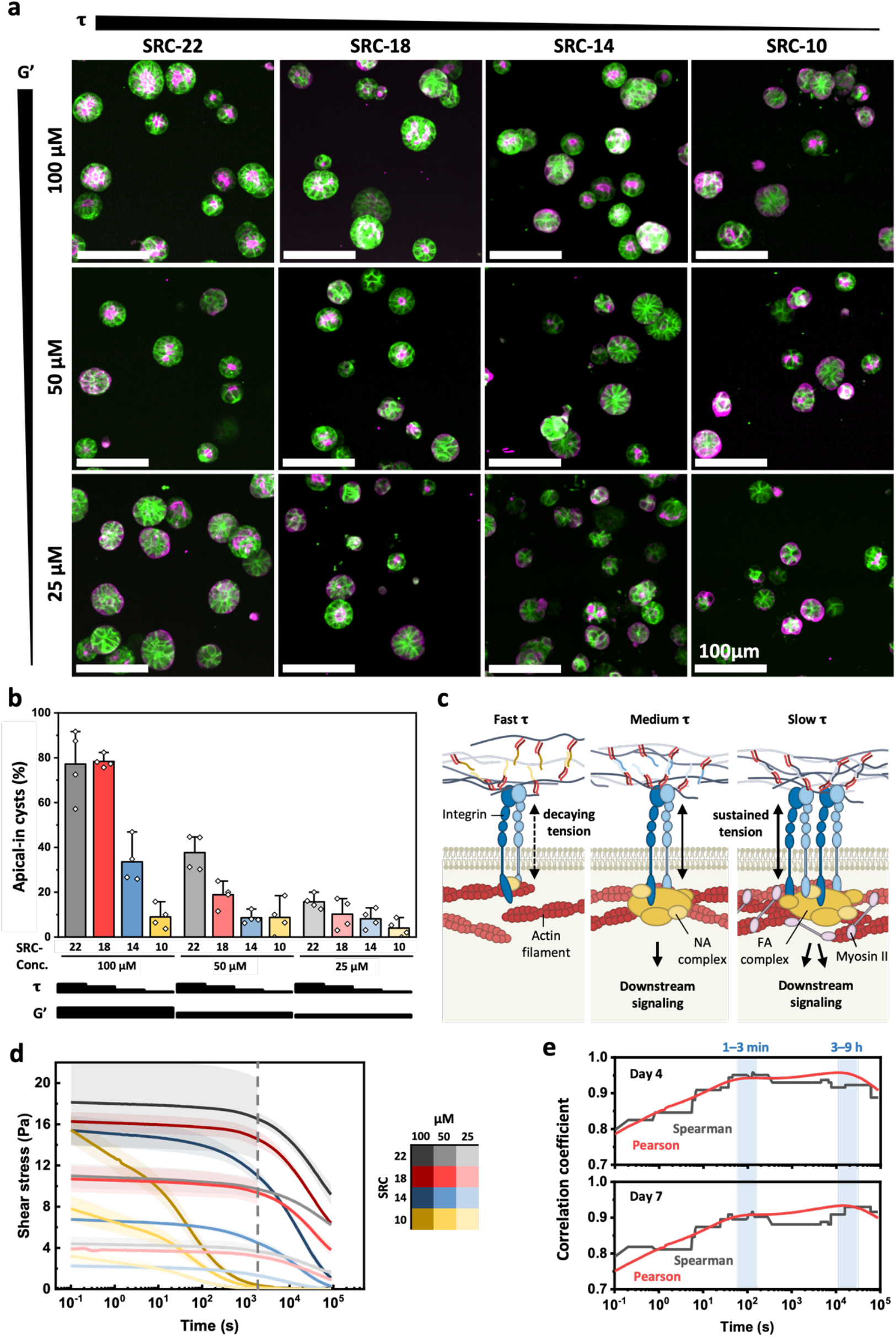
Systematic tuning of *DyNAtrix*’ viscoelasticity reveals two critical time scales for mechano-signaling processes involved in cell polarization. **a**, Confocal images of MDCK cyst (day 4) cultured in *DyNAtrix* with different stiffness and stress relaxation (see also Supplementary Figures S6–8.) **b**, Fraction of apical-in cysts cultured in *DyNAtrix* with different stiffness and stress relaxation combinations. **c**, Scheme of the tension between matrix and cells. (NA: nascent adhesion; FA: focal adhesion) **d**, Stress-relaxation curves of each of the 12 *DyNAtrix* variants. For each condition, the data is displayed as the average and standard deviation obtained from three independent step-strain experiments for data below 2000 seconds, and two independent step-strain experiments for data beyond 2000 seconds. The long-term relaxation data was normalized to the short-term data, and the transition between the two time domains is indicated by the dashed line. **e**, Correlation of the fraction of apical-in cysts on days 4 and 7 with the time-dependent shear stress obtained from the step-strain experiments. The time scales of two peaks in the time windows 1–3 min and 3–9 h are highlighted.

The effects of stiffness and stress-relaxation can be unified by considering the effective mechanical tension that builds up between the cells and the surrounding matrix (Figure 3c–e). The active contraction of the cytoskeleton creates tensile forces that induce conformational changes in integrins and adaptor proteins like talins. These changes trigger further protein assembly and ultimately modulate gene expression. In an ideal elastic material (like most synthetic matrices), these forces are proportional to the material’s stiffness. Correspondingly, previous studies on synthetic polymer gels had already revealed clear effects of matrix stiffness on differentiation and morphogenesis in epithelial cells^45^. However, in living tissues, which are viscoelastic, the local mechanical tension decays at a rate that depends on the ECM’s stress-relaxation behavior.

Crucially, cellular mechano-sensing not only depends on whether the critical mechanical tension builds up, but also whether it can be sustained on the timescale required for intracellular signal transduction to occur (Figure 3c). If so, integrins are activated and initially trigger the assembly of short-lived nascent adhesions (NA) that lead to downstream signaling. Longer timescales are required for the recruitment of myosin, which is needed for the maturation and reinforcement of the larger and more stable focal adhesion (FA) complexes.

But what timescale is relevant for the decision-making processes involved in the polarization of MDCK cysts? We hypothesized that our material platform provides a unique opportunity to answer this question by correlating systematic changes in its viscoelasticity with morphological changes. Non-normalized stress-relaxation profiles of the 12 different *DyNAtrix* gels provided shear–stress data over a wide range of timescales (Figure 3d). Each curve predicts the decay of tension along the matrix–integrin–cytoskeleton axis in a cell culture experiment. The fraction of apical-in cysts was therefore expected to be correlated with the non-normalized shear stress values measured at different time points. Based on this concept, we calculated *Pearson’s* correlation coefficient (r) and *Spearman’s* rank correlation coefficient (ρ) for every single time point of the stress-relaxation profiles (Figure 3e, see Supplementary Notes for details). Spearman coefficients reached very high values with a maximum between 1–3 minutes (ρ = 0.92–0.95). Pearson’s correlation coefficient plot exhibited a shoulder at the same time scale, reaching similarly high values (r = 0.91–0.94). Both correlation coefficients drop notably with decreasing timescale, indicating that cell polarization relies on mechanical cues that need to be sustained for more than a few seconds. Both Pearson and Spearman plots retained very high correlation coefficients on longer timescales, hinting at a second maximum in the 3–9 hour time range, which was particularly pronounced for day 7 cysts (r = 0.95, ρ = 0.95).

The two distinct “resonance” bands in the correlation plots are observed in all repeat experiments, both when correlating individual biological repeats to the average of the stress-relaxation data (4 repeats, Supplementary Figure S9 a-b) and when correlating individual stress-relaxation repeats with average experimental cyst polarity data (3 repeats, Supplementary Figure S9 c-d). The position of these bands can be linked to well-characterized mechano-biological processes known to occur in similar temporal windows. Firstly, the band at 1–3 minutes can be explained by mechanical forces and loading rates for integrin–ligand binding, which were characterized in several recent single-molecule studies^46–48^. Jo et al. reported loading rates of ∼1 pN/s for epithelial cells^46^. This value implies that it takes approximately 40 seconds to reach integrin’s peak tension of 40 pN (ref. 46), explaining why mechanical properties characterized on time scales much shorter than 40 seconds (or measured at oscillation frequencies significantly exceeding 0.025 Hz) would correlate poorly with cell polarization. Importantly, the lifetimes of the integrin–ligand bonds and of nascent adhesions are on the order of 60–100 seconds^49–51^. This transient nature provides an explanation for why the correlation trend stagnates for time scales beyond a few minutes. Thus, this first band (1–3 min) likely reflects the cell’s immediate mechano-sensory machinery.

We surmise that the second band between 3–9 hours corresponds to one or more mechano-signaling processes that require prolonged mechanical adaptation, including changes in protein synthesis and cytoskeletal reinforcement. As described above, nascent adhesions can mature into focal adhesions and fibrillar adhesions. This process can persist for up to several hours^49^ and requires the crosslinking of F-actin by myosin II^52^. Indeed, enrichment of myosin at the cell–matrix interface is pronounced in *DyNAtrix* with τ = 11 h and 1.1 d, but absent in *DyNAtrix* with 1 = 30 s (Figure 2d). The high correlation of sustained mechanical tension in the 3–9 hour time window with MDCK cyst polarization is therefore consistent with the timescale of adhesion complex maturation.

Intriguingly, the existence of different mechano-signaling processes taking place on well-separated time scales implies that it can be possible to selectively activate one pathway (but not the other) by adjusting the material’s stress relaxation time to an intermediate value between the two bands (i.e., between 3 min and 2 h). This idea is supported by the existence of exotic type III cysts in *DyNAtrix* if and only if it is set to the intermediate relaxation time of τ = 75 minutes. Under this specific condition, nascent adhesions and cytoskeletal responses take place, but the slower process of basal myosin enrichment is inhibited (Figure 2d,e). Thus, *DyNAtrix* makes it possible to isolate the effects of different signaling pathways without genetic modifications to the cells.

### Reversible τ-switching allows real-time guidance of cyst development

SRCs allow the programming of diverse cell-instructive mechanical properties in *DyNAtrix*. This helps to mimic some of the characteristics of living tissues and to interrogate mechano-signaling processes. However, living tissues are not static materials; they are subject to dynamic regulation, continuous remodeling, and changes in their mechanical characteristics throughout an organism’s development. We therefore sought to implement a mechanism by which *DyNAtrix*’ viscoelasticity could not only be adjusted during the experiment’s setup but also dynamically adapted in *ongoing* tissue culture to ideally support and actively guide cell development. To this end, we upgraded *DyNAtrix* with a new type of *switchable* stress-relaxation crosslinker (sSRC) (Figure 4a). The sSRC contained an 18-nt overlap domain, as well as an additional single-stranded domain located upstream of the overlap domain. This single-stranded domain served as a recognition site to capture a specific DNA signal, called the *fragilizer*. The *fragilizer* was designed to first bind the recognition sequence and then displace up to 6 bases in the overlap domain. This dynamic invasion of the overlap domain effectively shortens its length from 18 to 12 nucleotides. The *fragilizer* contains a single-stranded overhang (“toehold”) domain that allows its removal by adding a second strand, called the *reinforcer*. The system undergoe a toehold-mediated strand displacement (TMSD)^53–55^ reaction upon addition of the *reinforcer*. In this reaction, the *reinforcer* first binds to and then separates the *fragilizer* from the crosslinker, yielding a waste product called *fragiforcer*. As a result, the original crosslink is restored to its initial state. The *fragiforcer* can diffuse out of the hydrogel and be removed in subsequent medium exchanges. This specific design of sSRC, fragilizer, and reinforcer is termed **sSRC-18-12**.

**Figure 4.**
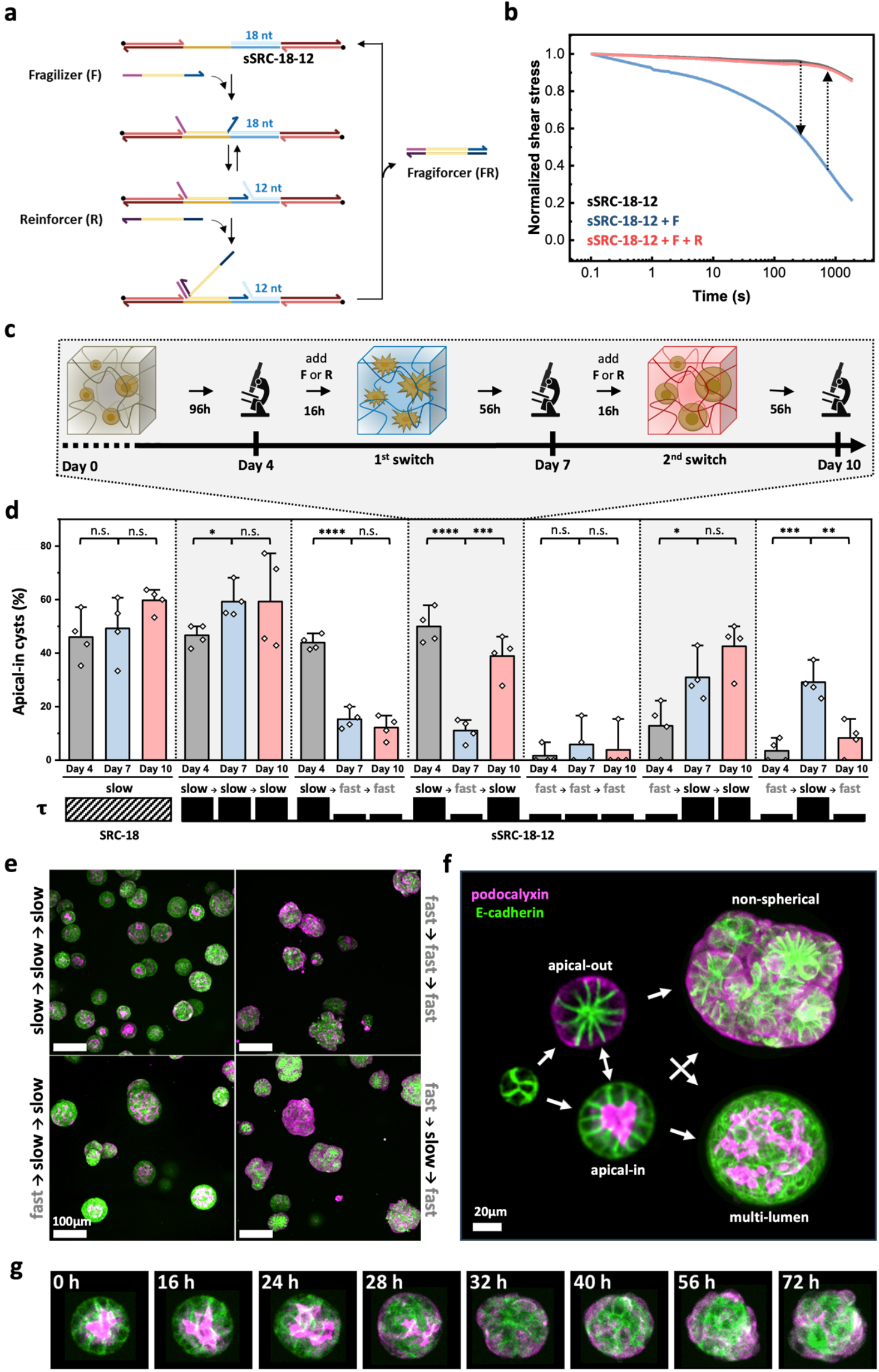
Control over MDCK cyst polarity via dynamic stress-relaxation switching in *DyNAtrix*. **a**, Scheme of **sSRC-18-12** and its switching mechanism. **b**, Normalized stress-relaxation curve of *DyNAtrix* crosslinked with the sSRC at the initial stage (dark grey), with fragilizer included (blue), and after successive addition of fragilizer and reinforcer (red). **c**, Scheme of *in situ* stress–relaxation switching during cell culture. **d**, Fraction of apical-in cysts cultured in *DyNAtrix*, responding to different temporal patterns in the material’s stress relaxation. Statistical analysis was performed using two-sample t-test (assuming equal variances). n.s., not significant; *, P < 0.05; **, P < 0.01; ***, P < 0.001; ****, P < 0.0001. **e**, Representative confocal microscopy images of MDCK cysts cultured with different stress-relaxation pathways (imaged on day 10). Extended data is shown in Supplementary Figure S12. **f**, Illustrative confocal microscope images of cysts exhibiting different morphologies depending on the cyst’s developmental stage and *DyNAtrix’* stress-relaxation history. From left to right, the images show an early unpolarized cyst (day 3), polarized cysts (day 6), and later cysts with complex multi-lumen and non-spherical morphologies (day 10). **g**, Time-lapse development of a single cyst initially grown in slow-relaxing *DyNAtrix*, imaged immediately after switching to fast stress-relaxation on day 4. Time stamps indicate the time elapsed after adding the fragilizer. Time-lapse movies of polarity inversion after τ-switching are shown in Supplementary Videos 1 and 2.

The proposed mechanism was validated in **sSRC-18-12**-crosslinked *DyNAtrix* via step-strain measurements (Figure 4b). In the absence of an additional signal strand, the stress-relaxation profile of **sSRC-18-12** closely matched that of the non-switchable **SRC-18** (Supplementary Figure S10). When *fragilizer* (2 eq.) was added to *DyNAtrix*, τ decreased from several hours to 7 minutes. This new value lies well between those of **SRC-10** (τ ∼30 s) and **SRC-14** (τ ∼75 min), confirming that stress-relaxation is predictably modulated through strand-invasion of the *fragilizer* into the overlap domain. Finally, when *reinforcer* (4 eq.) was added to the *fragilizer*-modified *DyNAtrix* gel, the original stress-relaxation profile was restored, confirming that the stress-relaxation regulation mechanism was fully reversible.

We examined the impact of dynamic stress-relaxation changes on apical-basal polarization at different time points of cyst development (Figure 4c–g). MDCK cells were embedded in *DyNAtrix* crosslinked by 100 µM **sSRC-18-12** and cultured for a total of 10 days. The cells were imaged on days 4, 7, and 10. During this time, *fragilizer* and *reinforcer* signals were added to the medium to switch between *fast* (7 min) and *slow* (11 h) stress relaxation. We chose three time points for triggering this transformation: on day 0 (i.e. as part of the cell culture setup), on day 4, and on day 7 (Figure 4c, Supplementary Figure S11). We studied the effects of six different switching patterns on cyst development (e.g., slow⟶fast⟶slow, fast⟶slow⟶slow, etc.) (Figure 4d,e; Supplementary Figure S12). Intriguingly, cyst polarity responded reliably to the temporal patterns in stress-relaxation changes. In control experiments (slow⟶slow⟶slow, fast⟶fast⟶fast) the average cyst polarity remained unchanged and was indistinguishable from a non-switchable reference *DyNAtrix*. However, switching *DyNAtrix* in the pattern slow⟶fast⟶slow triggered the inversion of apical-in cysts to apical-out cysts and back to apical-in cysts. Conversely, the fast⟶slow⟶fast pattern triggered apical-out-cysts to form apical-in cysts, which again inverted to apical-out cysts. All other tested switching patterns caused robust corresponding changes in cyst polarity.

To reveal the dynamics of polarity inversion, we imaged apical-in MDCK cysts in response to the stress-relaxation switching over 3-days, with a time interval of 2 hours (Figure 4g, Supplementary Videos 1 and 2). Upon addition of fragilizer to the culture, the cysts’ lumen began deforming gradually over about one day, yet no progressive accumulation of podocalyxin was detected on the basal side. Instead, the ultimate switch from apical-in to apical-out morphology took place rapidly within the timescale of less than 2 hours.

Besides polarity inversion, we observed several additional effects related to the switching of matrix stress relaxation: (1) when grown under changing stress-relaxation conditions, cysts were more prone to developing non-trivial (non-spherical or multi-lumen) morphologies. Thus, polarity inversion can provide access to more exotic multicellular structures. (2) Cysts grew larger when they had experienced at least one time segment with fast stress relaxation (Supplementary Figure S13a). A time window of fast stress-relaxation might aid cyst expansion into the matrix, while also allowing mechano-signaling to occur once switched to slow relaxation. (3) The material’s relaxation time during the final culture stage determined whether cysts were predominantly spherical or developed into non-spherical morphologies (Supplementary Figure S13b). We suspect that a slow relaxing environment supports the build-up of larger contractile forces that restore the rounded shape of the cyst, while fast stress relaxation leads to weaker contractile forces and thus a less spherical shape. A collage of the different morphologies and developmental pathways is shown in Figure 4f. Overall, our experiments demonstrate that *in situ* switching of stress relaxation in *DyNAtrix* presents a powerful tool for studying and directing cellular morphogenesis.

## Discussion

This study demonstrates a new approach for probing and guiding mechano-biological processes in 3D cell culture. Dynamic DNA crosslinks allow the decoupling of matrix stiffness and stress-relaxation characteristics, providing insights into mechano-signaling processes underlying apical-basal polarization in MDCK cells. The evolved design of *DyNAtrix* offers an expanded range of relaxation times (from seconds to days) and a dynamic mechanism to change the material’s stress relaxation characteristics mid-experiment.

Our novel correlation analysis (see Supplementary Notes) provides a straightforward methodology to associate one or more *time-resolved mechanical properties* (e.g., the stress remaining at each time point in a step-strain experiment) with one or more *biological outcomes* (e.g., the fraction of apical-in cysts). In this way, one can identify the *timescale* that is most relevant for the development of an observed phenotype, shedding light on the rate of the underlying mechano-biological processes. Using this method, we find that two time windows are particularly relevant for mechano-signaling processes influencing MDCK cell polarization in *DyNAtrix*: a rapid response on the order of 1–3 minutes, which is linked to integrin-mediated adhesion dynamics; and a much slower process occurring over 3–9 hours, which coincides with cytoskeletal reinforcements and focal adhesion maturation.

All previous biophysical analyses report “matrix stiffness” as the plateau modulus, typically measured at ∼1 Hz. However, our study demonstrates that this frequency can have limited physiological relevance for cellular mechano-sensing. Given the timescale of integrin force loading rates and nascent adhesion lifetimes, a measurement timescale of at least 40 seconds (or ≤ 0.025 Hz in oscillatory rheology) is more meaningful for matrix stiffness. Since no single stiffness value can capture the full complexity of ECM mechanics, it is crucial to report mechanical matrix properties over a wide range of timescales.

The need for controlling cell polarization is crucial for addressing a wide range of questions in developmental biology and advanced organoid research^56,57^. To the best of our knowledge, this work is the first systematic study showing how the polarization of epithelial cells can be dynamically changed simply by altering time-dependent ECM mechanics, without changing biochemical signals or medium composition. Slow stress relaxation facilitated lumen formation and enhanced myosin enrichment at the cell–matrix interface, highlighting the importance of mechanical tension in driving cellular organization. Our findings provide further support for the growing body of research demonstrating that seemingly subtle aspects of ECM mechanical characteristics can have profound effects on cell development^17^. Interestingly, a recent study by Ruiter et al. found that kidney organoids embedded in matrices with two different stress relaxation times (4,100 s and 39,000 s) also formed different morphologies, showing subtle differences in polarity^58^. The exceptionally broad dynamic range of stress-relaxation times in *DyNAtrix* therefore promises to provide control over the development of various organoid models.

An unprecedented degree of *in situ* control is achieved through the switchable stress-relaxation crosslinker (sSRC), enabling real-time modulation of matrix viscoelasticity. The dynamic strand displacement reaction permits active regulation of cellular behavior during development, as demonstrated by the reversible polarity inversion and the induction of complex multicellular morphologies in MDCK cysts. Such capabilities open new avenues for tissue engineering, where guiding cell fate and structure through adaptive mechanical environments could significantly enhance the creation of biomimetic tissues. Inverting cell polarity is also relevant for disease modeling. For instance, metastasis of cancerous tumors frequently involves reversible changes in cell polarity^12^. Multicellular epithelial cysts cultured in basement membrane matrices have the apical side of the cell structure facing the enclosed lumen (apical-in). In some cases, this apical configuration can hinder *in vitro* research, as the apical side is difficult to access for controlled exposure to nutrients, toxins, or pathogens^57^. Microinjection is a traditional method for delivering these targets into the lumen; however, it is time-consuming and can damage the organoids. The reversible polarity inversion in *DyNAtrix* might therefore offer a simple, non-invasive, and high-throughput alternative to microinjection.

Overall, the reported methodology advances our understanding of mechano-transduction in epithelial cells and serves as a versatile platform for investigating dynamic tissue mechanics. This innovation bridges the gap between static synthetic matrices and the complex, adaptable nature of living tissues, paving the way for advanced studies in cell biology, disease modeling, and regenerative medicine. Future studies should explore the use of *DyNAtrix* for more complex cell systems and organoid models to further validate its potential for basic research and clinical applications. Currently, *DyNAtrix* is limited to elastic moduli below 500 Pa, making it most suitable for soft tissues like those found in the brain, gut, liver, kidney, and placenta. To broaden its applicability, future developments will focus on accessing higher stiffness regimes in the range of 0.5–20 kPa. Moreover, alternative adhesion ligands will be implemented to study the specific individual effects of mechanical cues transmitted through different receptor–ligand bonds.

## Methods

### Materials

Solvents and reagents were purchased from commercial sources and used as received, unless otherwise specified. Methanol (ACS reagent grade) was obtained from Fisher Scientific. Molecular biology grade acrylamide (catalog number A9099), sodium acrylate (catalog number 408220) and ammonium persulfate (APS; catalog number A3678) were purchased from Sigma-Aldrich. Ultrapure N,N,N′,N′-tetramethylethylenediamine (TEMED; catalog number 15524010) was obtained from Thermo Scientific. Desalted oligonucleotides were purchased from Integrated DNA Technologies (IDT). Amino acids for peptide synthesis were purchased from IRIS Biotech GmbH. Nitrogen gas (>99.999%) was used under inert conditions and supplied by an in-house gas generator. To ensure an inert condition, it was purified through a Model 1000 oxygen trap from Sigma-Aldrich (catalog number Z290246). Reagents with unreacted acrylamide groups were stored at 4 °C or −20 °C, protected from unnecessary exposure to light.

### Peptide synthesis

RGD peptides (sequence: G(acryl-K)GGGRGDSP) were synthesized on a Liberty Blue HT12™ automatic and microwave-assisted peptide synthesizer (CEM GmbH) using a Rink Amide resin and a 9-fluorenylmethoxycarbonyl protection strategy. Amino acid activation was achieved by N, N’-diisopropyl carbodiimide, and ethyl cyanohydroxyiminoacetate. Acetylation of the N-terminus was performed via incubating the resin-bound peptides in acetic anhydride for 2 h. The reaction mixture was continuously stirred and vigorously saturated by bubbled nitrogen to prevent oxidation reactions and afterward washed three times with N, N-dimethylformamide. Deprotection of the amino acid side chains and cleavage from the resin was accomplished using a mixture of trifluoroacetic acid (φi = 87.5%), phenol (φi = 5%), triisopropylsilane (φi = 2.5%) and MilliQ-water (φi = 5%) for 3 h at room temperature. The crude peptide was precipitated in anhydrous diethyl ether, collected by vacuum filtration, and dried under nitrogen flow. Further purification of the peptides was realized by high-performance liquid chromatography (Agilent 1200, Agilent Technologies) on a preparative C18 column (10 μm particle size, 100 Å pore size, 250 × 30 mm, Phenomenex Ltd). A linear gradient of MilliQ-water/acetonitrile and trifluoroacetic acid (φi = 0.1%) was used as the mobile phase. Finally, the concentrated peptide-containing solution was lyophilized (Alpha 24 LD plus freeze-dryer, Martin Christ) and the obtained product was stored at -20°C afterward until further usage.

### Polymer synthesis

The polymer synthesis protocol is based on refs. ^36^ and ^37^. In brief, acrylamide (50 mg ml^τ1^), sodium acrylate (0.5 mg ml^τ1^) and acrylamide-labeled anchor strand DNA (1 mM, Supplementary Table S1) were co-polymerized in TBE buffer (100 mM Tris, 100 mM boric acid, 2 mM EDTA, pH 8.2). For the RGD-functionalized derivative, **P**^**+RGD**^, acrylated RGD peptide (sequence, G(acryl-K)GGGRGDSP) was added to the above solution (10 mM, final concentration). The polymerization was initiated by adding TEMED (0.025 wt%, final concentration) and APS (0.025 wt%, final concentration). To achieve high molecular weight and narrow size distribution, the reaction was carried out under high-purity nitrogen gas, which was passed through an oxygen trap on-site. The reaction was allowed to proceed for 24h, producing a highly viscous solution. NMR spectroscopy was used to assess the conversion^36^. For purification, the solution was diluted in 9x volumes of Milli-Q water and subsequently purified via precipitation with methanol. The pellet was re-suspended in Milli-Q water and stored in aliquots (∼2.5 wt%) at τ20 °C. NMR indicated reaction conversions of 98.4% and 98.8% for **P**^**+RGD**^ and **P**, respectively (Supplementary Figure S1). Analytical DNA binding efficiency tests confirmed that >90% of acrylamide-labeled DNA strands in the reaction had been incorporated as anchor strands into the polymer backbone and were accessible for binding complementary DNA strands (Supplementary Figure S2).

### Nuclear magnetic resonance (NMR) spectroscopy

NMR samples were prepared by diluting 100 μL of the unpurified product in 700 μL of D_2_O. ^1^H NMR spectra were recorded at 30–32 °C on a 500 MHz spectrometer (Bruker) with a 2-second acquisition time and 32 transients. Chemical shifts (δ) are reported in parts per million (ppm) downfield from tetramethyl silane (TMS). The ^1^H NMR shifts are referenced to the residual hydrogen peak of D_2_O (4.79 ppm).

### Polyacrylamide gel electrophoresis (PAGE)

The binding capacity of **P**^**+RGD**^ and **P** was determined via PAGE. A native PAGE gel was prepared using 40 % (w/w) acrylamide/bis-acrylamide (19:1) stock. The samples were prepared in 1x TE buffer with a final concentration of 150 mM NaCl and 0.01 % (w/v) solutions of **P**^**+RGD**^ or **P** (2 μM anchor strands). The complementary strand of the anchor strand (Strand ID#1) was added at series equivalent from 0.2x∼ 2x. The samples were annealed on a thermal cycler at 95 °C for 2 minutes, followed by instant cooling to 4 °C and holding for 3 minutes. The annealed products were loaded onto a 10 % native polyacrylamide gel and run at 120 V on a Mini-Cell® Novex system from Life Technologies in 0.5x TBE buffer via a Consort EV265 power supply. The gel was stained with 1x SYBR™ Gold for 15 minutes and scanned on Typhoon FLA 9000 Scanners (GE Healthcare Life Sciences) via a blue LD laser (excitation at 473 nm) at 25 μm/pixel resolution. Image analysis and densitometric quantification of the gel bands were performed in Fiji.

### *DyNAtrix* preparation

The protocol is based on ref. ^36^. In brief, precursors A and B were prepared. A contains the reverse splint of SRCs and the corresponding blocking strands; B contains the DNA-functionalized polymer, the forward splint of SRCs and the corresponding blocking strands. In both A and B, the blocking strands are always 2 folds of the SRCs. By adjusting with the 10X DMEM concentrate, the precursors were finally in 1X DMEM. Next, the precursors were annealed by heating up to 70°C and cooling down to 4°C with the rate of -2°C min^-1^. Then, they were stored at 4°C until use. By mixing the two precursors and incubation at 37°C, a 1% (w/v) gel is formed with maximum 100 µM crosslinks, which resulted from 100 µM forward splints binding with 100 µM reverse splints of SRCs. In this study, we formed gels with 100, 50 and 25 µM crosslinks.

### Oscillatory rheological measurements

Viscoelasticity measurements were conducted using Anton Paar MCR301 rheometer with a 25-mm-diameter cone-plate geometry (cone angle 0.5°). Temperature control was ensured by installing a Peltier system PTD-200 (Anton Paar) within the measurement area. To maintain high humidity and prevent evaporation artifacts, a wet paper cylinder was placed around the plate’s circumference. Stress–relaxation properties were measured at 15% strain, where the strain was kept constant while recording shear stress over time. Frequency sweeps were conducted from 0.1 Hz to 100 Hz at 10% strain. The real-time gelation test was measured at 10% strain and 1.6 Hz. For the real-time gelation test, the temperature started and stayed at 4°C for 5 min, gradually elevated from 4 to 20°C within 3 min, then from 20 to 37°C within 30 min, and stayed at 37°C for 240 min. All *DyNAtrix* samples were injected onto the rheometer after freshly mixed the precursors. For the validation of switchable stress relaxation, the precursors were prepared by including the fragilizer (F) in precursor A and the reinforce (R) in precursor B. The concentration ratio of sSRC:F:R was 1:2:4.

### MDCK cell culture

Two types of auto-fluorescent MDCK cells (strain II) were provided by Alf Honigmann’s group. One of them expresses E-cadherin-mNeonGreen and podocalyxin-mScarlet; the other expresses myosin-mNeonGreen and mucin-Halo. The cells were cultured in minimum essential medium (MEM) with GlutaMAX™ Supplement (Gibco, catalog number 11140050), 1%MEM Non-Essential Amino Acids Solution (100X) (Gibco, catalog number 41090028), 1mM Sodium Pyruvate (Gibco, catalog number 11360070), 5% FBS (Sigma Aldrich, catalog number F7524), and 100U Penicillin and 0.1 mg Streptomycin (Sigma Aldrich, catalog number P4333) at 37°C and 5% CO2 in a humidified incubator. The cells were grown until 80%-90% confluence in the T25 flask before detachment with 0.25% trypsin-EDTA solution.

### 3D cell culture

After detachment, the MDCK suspension was centrifuged, and the cell pellet was resuspended with UltraMDCK™ Serum-free Renal Cell Medium (Lonza, catalog number BEBP12-749Q). This was repeated two times to move the serum completely. Then the suspension was mixed with the precursors on ice. The final cell density in the gel was 4 × 10^5^ cells/mL. Next, the mixture (5 μl per micro-well) was placed in the micro-well inserts (*ibidi GmbH*, catalog number 80409) attached in 6-well plates. The samples were incubated at 37 °C for 60 min to trigger heat-activated gelation. After gelation, 5mL of UltraMDCK™ Serum-free Renal Cell Medium was added to each well (the whole insert was covered by medium).

For the static tuning experiment, we covered the 5 μl *DyNAtrix* with 10 μl starPEG-Heparin gel to prevent the gel from swelling. This measure ensured the constant concentration of cell adhesion peptides and other gel components. The thiolated 4-arm-poly(ethylene glycol) (JenKem, A7008-10) and the maleimide-functionalized heparin (molecular weight: 15000, 6 maleimide groups per heparin) were kindly provided by Carsten Werner’s lab. The starPEG-Heparin gel was formed with the final concentration of 10.6 mg/mL thiolated 4-arm-poly(ethylene glycol) and 15 mg/mL maleimide-functionalized heparin in phosphate-buffered saline (PBS).

### *In situ* stress relaxation switching

The switching was done by adding displacement strands, fragilizer and reinforce (Supplementary Table S1). After imaging on day 4, most of the medium was removed. Only around 200 μl of the medium was kept in the inserts. 25 μl of the 2M displacement strands were added into the insert and gently mixed with the remaining medium. Here, the concentration of the displacement strands was 200 μM, which was two folds of the crosslinker concentration. After overnight incubation (around 16h), the medium was refilled. On day 7, the samples were imaged and the 2^nd^ switch proceeded as day 4 (Supplementary Figure S11).

### Confocal microscopy

Images were acquired on an Andor Dragonfly Confocal Microscope (Oxford Instrument).

## Acknowledgments

E.K. acknowledges funding by the Federal Ministry of Education and Research of Germany (BMBF) in the program NanoMatFutur (grant no. 13XP5098). The authors thank Isabell Jeglinski for synthesizing the modified RGD peptide and Dr. Hartmut Komber for assistance with NMR measurements.

## Author contributions

E.K. conceived the project. S.K.H. and E.K. designed the experiments. S.K.H. carried out polymer synthesis and characterization, DNA sequence design, oscillatory rheology, gelation, and cell culture experiments. A.H. developed the genetically modified MDCK cells. E.K. carried out the correlation analysis. C.W., A.H. and M.M. participated in planning the cell culture experiments and data analysis. S.K.H. and M.M. carried out some long-term live-cell imaging experiments. S.K.H. and E.K. wrote the initial manuscript draft. All authors discussed the results and helped revise the manuscript.

