## Supplementary Information for "Programmable viscoelastic hydrogels uncover mechano-sensing timescales and direct cell polarity"

### Supplementary Notes

#### Correlation of stress-relaxation data with MDCK cyst polarity

This section describes how we generated the correlation plots between the time-resolved stress-relaxation profiles of the different *DyNAtrix* variants and the measured MDCK cyst polarity. We first summarize the experimental data that was used in these plots, then we present the mathematical approach used to calculate the correlation coefficients as a function of time, and finally try to give an intuitive explanation for this analysis. A visual description of the process with example data is shown in Supplementary Note Figure 1.

#### *1. Input data*

By assembling **P$^{+RGD}$** with 4 different SRCs, each at 3 different concentrations, we produced 12 different gels exhibiting distinct viscoelastic properties. Two datasets were measured for each gel, one that reflects a key mechanical property (stress-relaxation behavior), and one that reflects a morphological outcome (cyst polarity).

##### Stress-relaxation curve

For each of the 12 gels, a step-strain experiment was performed on a rheometer. At $t$ = 0, we imposed a constant shear strain ($\gamma$ = 15%) and measured the decay in shear stress $\sigma(t)$ over time (from seconds to many hours or days, depending on the sample). The result for each gel $i \in \{1, 2, \ldots, 12\}$ is a function $\sigma_i(t)$, where $t$ spans a broad range of times. We used **non-normalized** stress data from the rheometer to reflect the absolute stress levels that cells experience.

##### MDCK cyst polarity

MDCK cells were cultured in each of the 12 *DyNAtrix* gels, and at specific culture days (i.e., days 4 and 7) we quantified the fraction $P_i$ of cysts exhibiting an apical-in morphology. Thus, each gel $i$ is associated with a single (scalar) cyst polarity value, $P_i$, measured independently on day 4 and day 7.

Our goal was to study how well the stress that remains in the gel after a given time $t$ (in a step-strain experiment) correlates with the cyst polarity observed in that gel.

### 2. *Constructing time-resolved stress vectors*

There are 12 stress-relaxation curves:

$$\sigma_1(t), \sigma_2(t), \ldots, \sigma_{12}(t)$$

For each fixed time point t, we can therefore form a 12-dimensional vector of shear-stress measurements:

$$S(t) = (\sigma_1(t), \sigma_2(t), \ldots, \sigma_{12}(t))$$

### 3. *Constructing cyst polarity vectors*

Similarly, the measured fraction of apical-in cysts for each of the 12 gels on day 4 or day 7 can be expressed as a 12-dimensional vector:

$$P = (P_1, P_2, \ldots, P_{12})$$

We generated two such vectors, one for day 4 ($P_{day\ 4}$) and one for day 7 ($P_{day\ 7}$).

### 4. *Correlating stress and polarity as a function of time*

To analyze how well the stress $\sigma_i(t)$ at each time t in the stress-relaxation profile correlated with the observed polarity $P_i$ , we computed classical correlation coefficients between the vectors $S(t)$ and $P$. For each discrete time point $t$, we calculated Pearson's correlation coefficient and Spearman's rank correlation coefficient.

#### Pearson's correlation coefficient

Pearson's correlation coefficient as a function of time, $r(t)$, was calculated as:

$$r(t) = \frac{\sum_{i=1}^{12} [\sigma_i(t) - \bar{\sigma}(t)][P_i - \bar{P}]}{\sqrt{\sum_{i=1}^{12} [\sigma_i(t) - \bar{\sigma}(t)]^2}\sqrt{\sum_{i=1}^{12} [P_i - \bar{P}]^2}}$$

where $\bar{\sigma}(t)$ is the average of all $\sigma_i(t)$ over i = 1, 2, …, 12; and $\bar{P}$ is the average of all $P_i$ over $i$ = 1, 2, …, 12.

#### Spearman's rank correlation coefficient

Spearman's rank correlation coefficient as a function of time, $\rho(t)$, was calculated as:

$$\rho(t) = 1 - \frac{6\sum_{i=1}^{12} \left(R_{\sigma_i(t)} - R_{P_i}\right)^2}{12(12^2 - 1)}$$

where $R_{\sigma_i(t)}$ and $R_{P_i}$ are the rank orders of $\sigma_i(t)$ and $P_i$, respectively, when sorted from smallest to largest.

We sampled the full range of time points $t$ across several orders of magnitude (seconds to hours). Both Pearson and Spearman correlation plots were generated for day 4 and for day 7 polarity data, as shown in Figure 3e.

### 5. *Interpreting the correlation plots*

Plotting $r(t)$ and $\rho(t)$ versus $t$ reveals which time windows in the stress-relaxation decay show the strongest correlation with the measured polarity outcomes. In other words, the values in the correlation plots indicate to what extent the residual stress that persists on a particular timescale (following a deformation) is relevant for the mechanosensitive processes governing MDCK cyst polarity (e.g., integrin bond lifetimes at short timescales and cytoskeletal remodeling at longer timescales).

### 6. *Intuitive explanation*

Imagine you are participating in a study where researchers hand you twelve different materials, one after another—a piece of playdough (1), a gummy bear (2), a piece of jelly (3), a rubber ball (4), a lump of tar (5), and so on—each with different mechanical characteristics. Your task is to rank these material by how “stiff” they feel to you. Your assessment of the materials’ stiffnesses can be quite subjective. For instance, if you are a person who relies on quick first impressions, you may have only briefly touched each material. In that case you might decide the playdough and the rubber ball feel equally stiff. But if you spend more time steadily squeezing them, you discover that the playdough gradually yields (relaxes), while the rubber ball remains firm; you would therefore rank the ball as stiffer than the dough. You write down your assessments of perceived “stiffnesses” for the researchers, for instance: 1: soft, 2: soft, 3: very soft, 4: hard, 5: medium, ….

Back in the lab, the researchers measure each material’s *time-dependent* stress-relaxation behavior with a rheometer: they apply a deformation and track how much stress the material retains or loses over seconds, minutes, hours, and days. Some materials lose stress quickly (like playdough), while others hold stress longer (like the rubber ball). They then compare your “stiffness” rankings to the rheometer’s readouts at different time points. This way, they obtain a curve of correlation coefficients spanning the entire stress-relaxation window. A maximum at, for example, 100 seconds means your rankings best match the stress each material retains at 100 seconds, suggesting you spent about that long to decide how stiff it felt. When they pool data from many participants, they may see multiple peaks in this correlation curve. Each peak marks a timescale that many people happen to use (consciously or unconsciously) when evaluating the materials’ stiffness.

We use the same idea in our correlation analysis of cell culture data in *DyNAtrix*. When cells pull on the matrix, the tension they feel depends on how quickly or slowly the matrix relaxes under that pull. Cells themselves have characteristic timescales for sensing and responding to force, for instance, the timescale of integrin binding or focal adhesion assembly. Because the tension across integrin–matrix bonds is directly proportional to the measured stress in a step-strain test, integrin-dependent processes like cell polarity end up correlating most strongly with the matrix’s stress at the time intervals over which the cells are actually applying and sensing force.

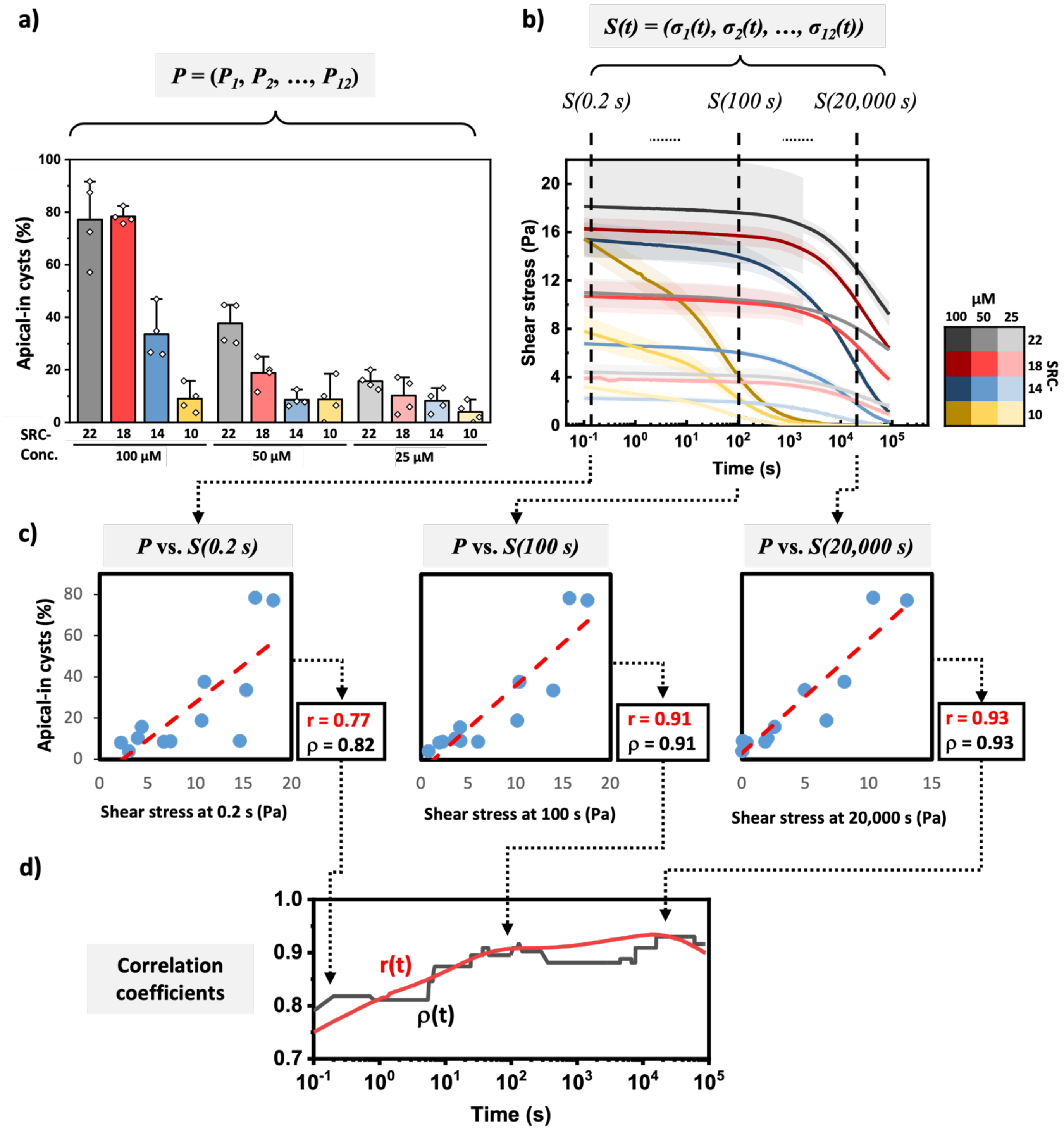


**Supplementary Note Figure 1. Correlation analysis. a)** The experimental data of the percentage of apical-in cysts in 12 different *DyNAtrix* variants is used to construct the vector *P*. Example for day-7 cysts: *P* = (77%, 78%, 34%, 9%, 37%, 19%, 9%, 9%, 16%, 10%, 8%, 4%). **b)** The time-dependent shear stress data from step strain experiments on the same 12 different *DyNAtrix* variants is used to construct a vectors for each time point in the stress-relaxation data, *S(t)*. Example for 100 second time point: *S(100 s)* = (18, 16, 15, 15, 11, 11, 7, 7, 4, 4, 2, 3). **c)** The correlation between the quantified polarities (vector P) and the experimental shear stresses in the corresponding *DyNAtrix* gels (vector S) is calculated for each time point of the step-strain data. Examples are shown for the 0.2 s, the 100 s, and the 20,000 second time points. The correlations yield a Pearson coefficient (r) and a Spearman rank coefficient (ρ) for each time point. **d)** Example plot of the timescale-dependent correlation coefficients for day-7 polarity and shear stress.

### Supplementary Figures

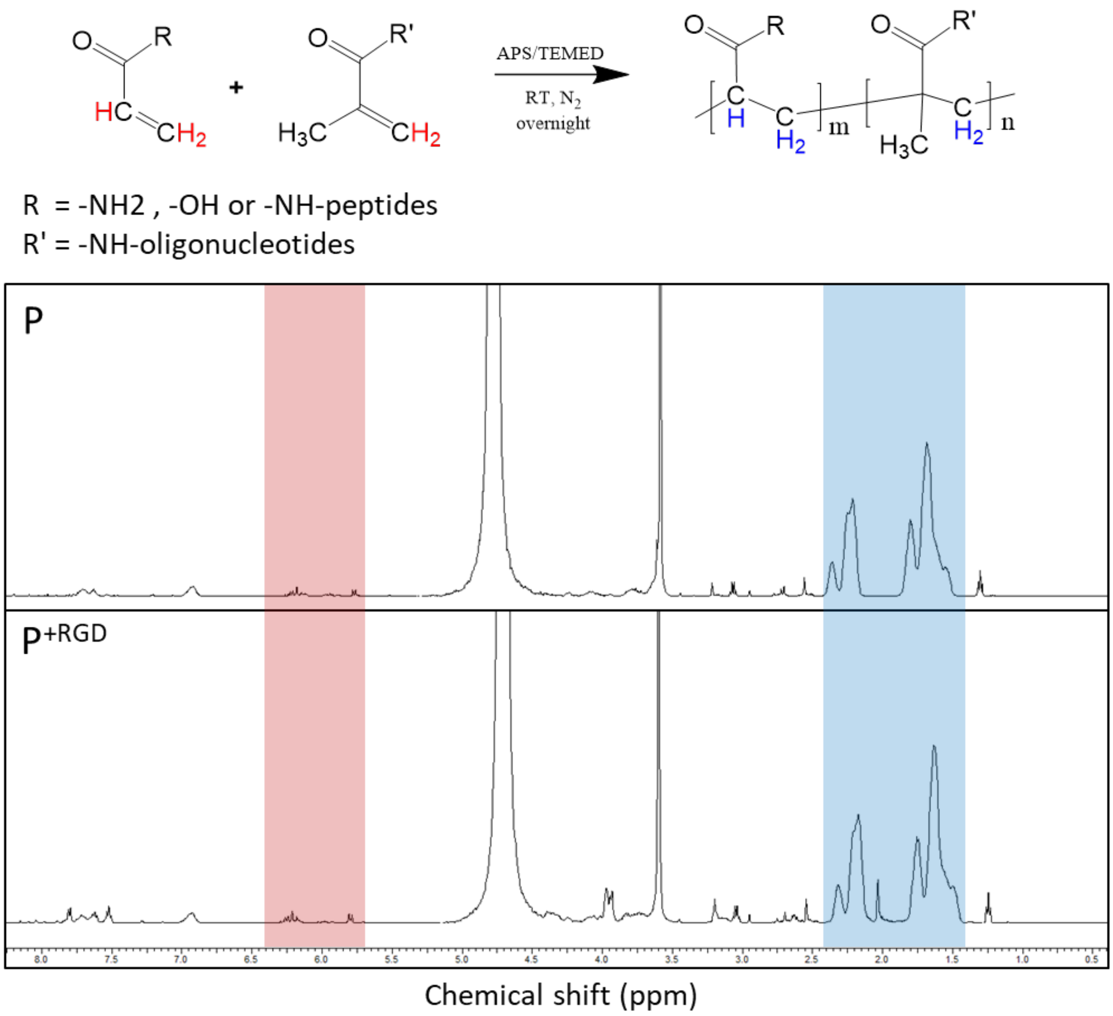


**Figure S1. $^1$H-NMR spectra of P and P$^{+RGD}$ in $D_2O$ after the reaction was completed and prior to methanol precipitation purification.** The conversion of the reaction was calculated by quantifying the free residual acrylamide monomer protons (δ~5.7--6.4; red region) vs. the polymer backbone protons (δ~1.4--2.4; blue region). The conversions are 98.4%, and 98.8% for **P** and **P$^{+RGD}$**, respectively. R = -$NH_2$, -OH or -NH-peptides; R' = -NH-oligonucleotides.

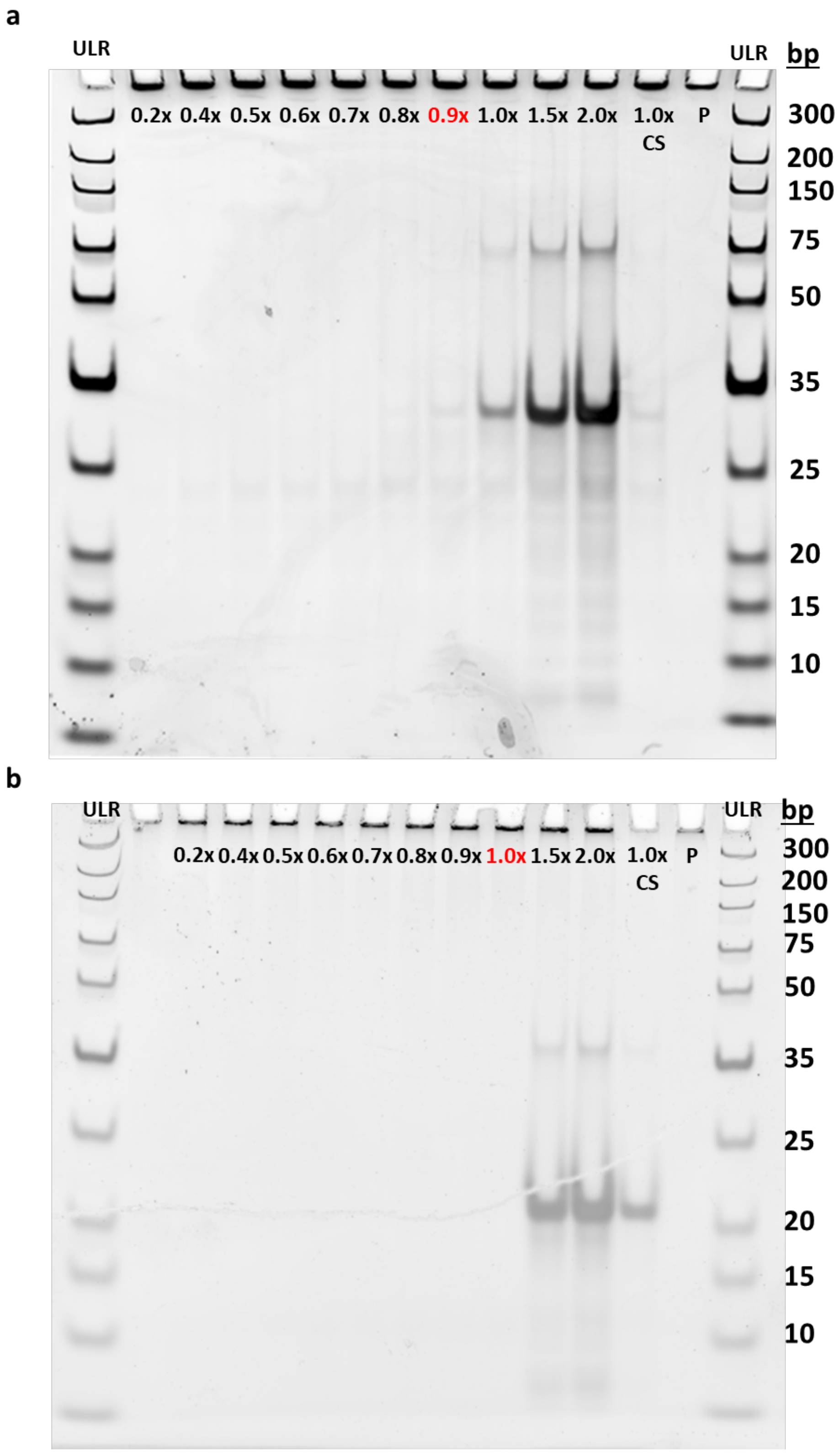


**Figure S2. DNA binding efficiency test to quantify the number of accessible anchor strands per polymer chain after purification. a, P. b, P$^{+RGD}$.** Take **P** as an example, 0.01% (w/v) polymer, which contains a theoretical concentration of 2 µM anchor strands, was mixed with the complementary strand (CS) at ratios ranging from 0.2x to 2.0x. Complementary strands at concentrations below 0.9x did not appear as distinct bands on the gel, indicating that they were quantitatively bound to the polymer. 1.0x of the complementary strands saturated the anchor strands and clearly revealed bands of excessive CS on the gel. Thus, the concentration of available anchor strands was determined to be approximately 1.8 µM. The results revealed that 90-100% of the anchor strands were incorporated with the polymer. (ULR: ultralow range DNA ladder; bp: base pair)

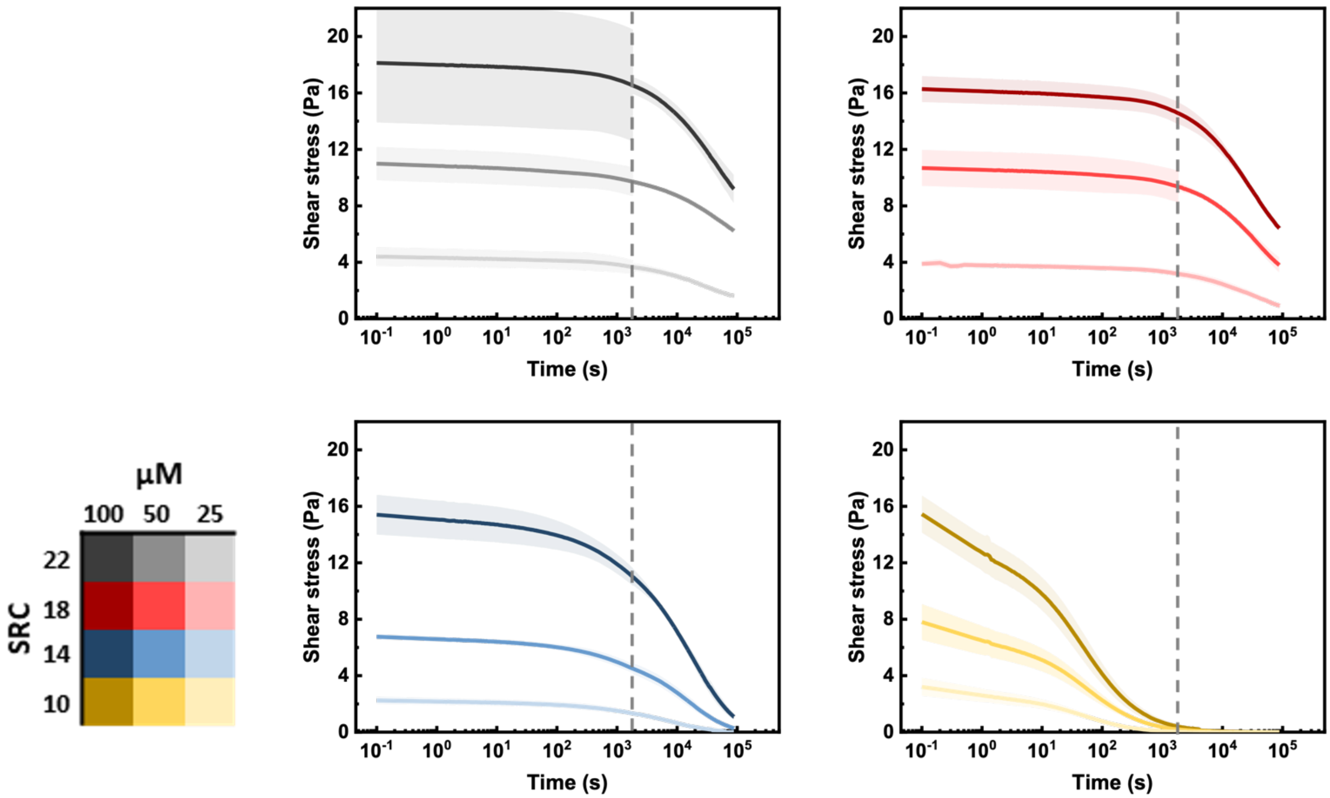


**Figure S3.** The stress-relaxation curves of the four non-switchable SRCs. For each condition, the data is displayed as the average and standard deviation obtained from three independent step-strain experiments times below 2000 seconds, and two independent step-strain experiments for data beyond 2000 seconds. The long-term relaxation data was normalized to the short-term data, and the transition between the two time domains is indicated by the dashed line.

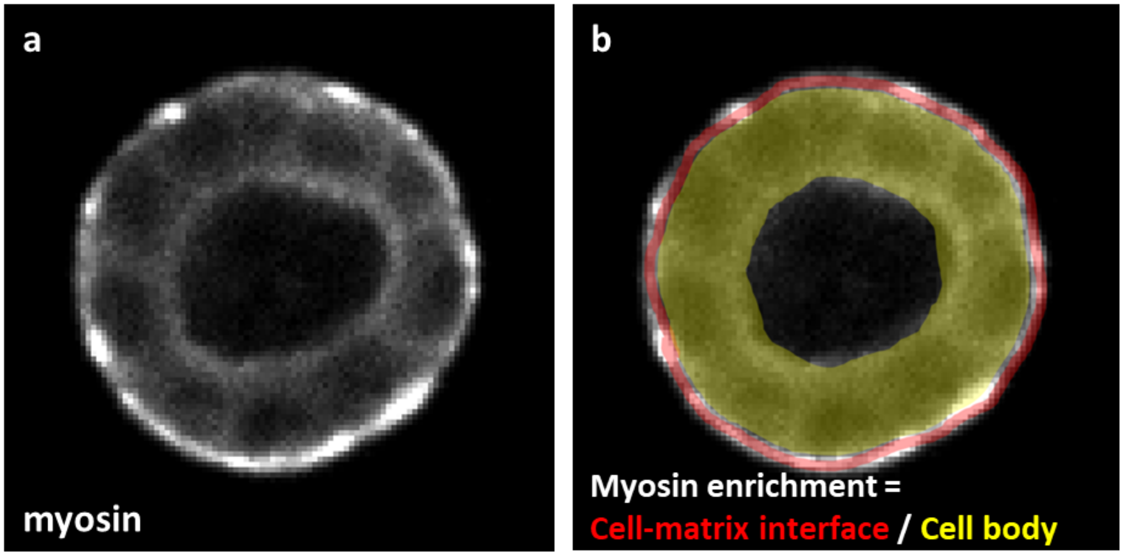


**Figure S4. Myosin enrichment analysis of MDCK cysts. a,** Confocal image of a cyst with myosin IIa labeled. **b,** Myosin enrichment is defined as the average intensity ratio of myosin at the cell-matrix interface and whole cyst. The intensity is defined as the average brightness of pixels in the highlighted regions.

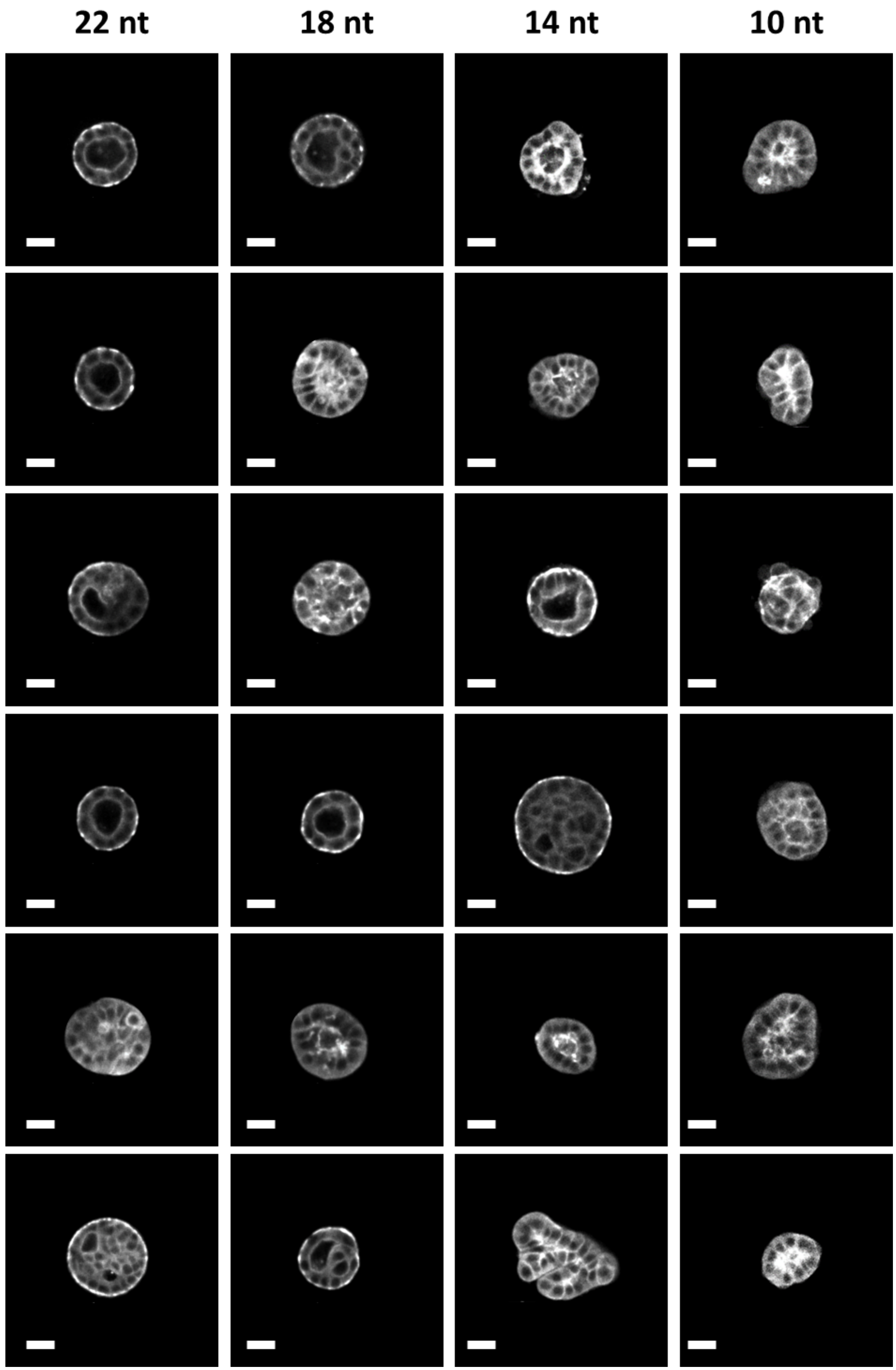


**Figure S5. Confocal images (cross-section) of 6 representative cysts for each gel condition.** Scale bar = 20 µm.

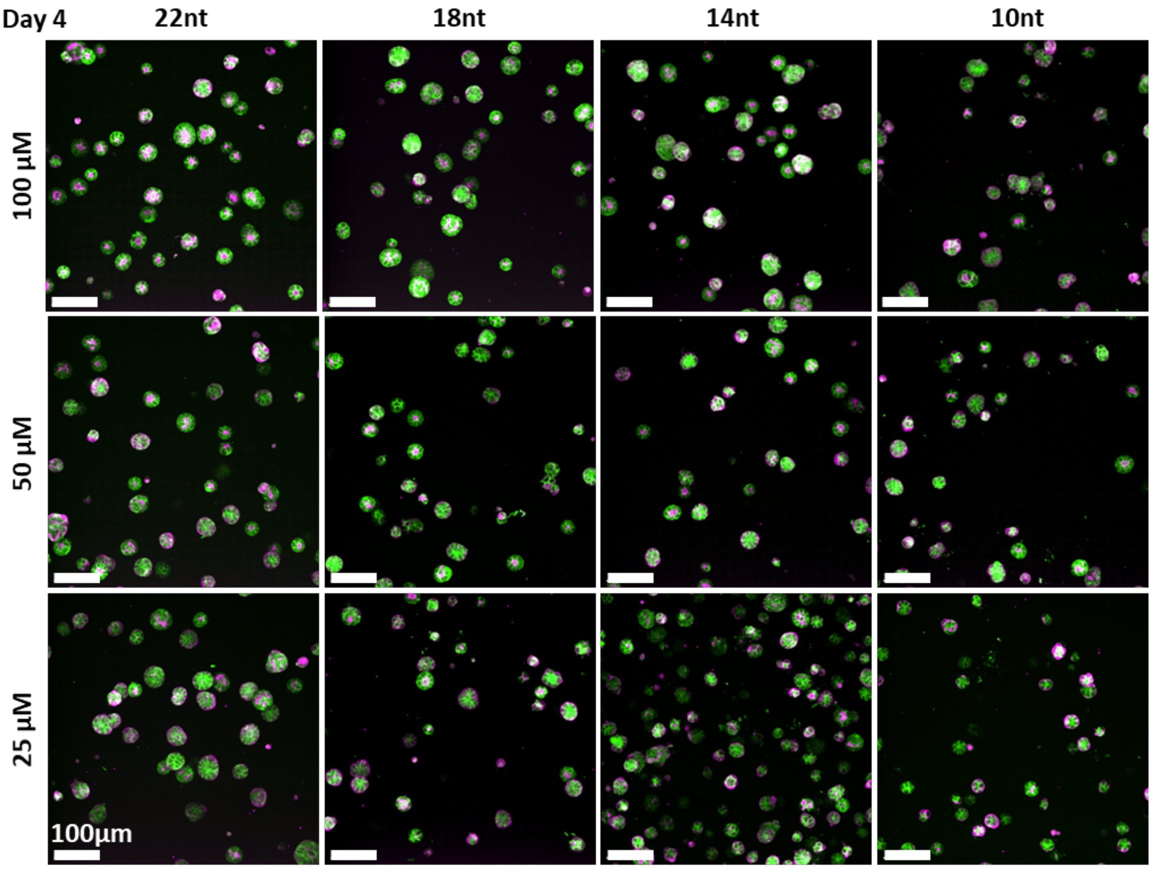

**Figure S6. Confocal images of MDCK cyst in *DyNAtrix* with different stiffness and stress relaxation (day 4).**

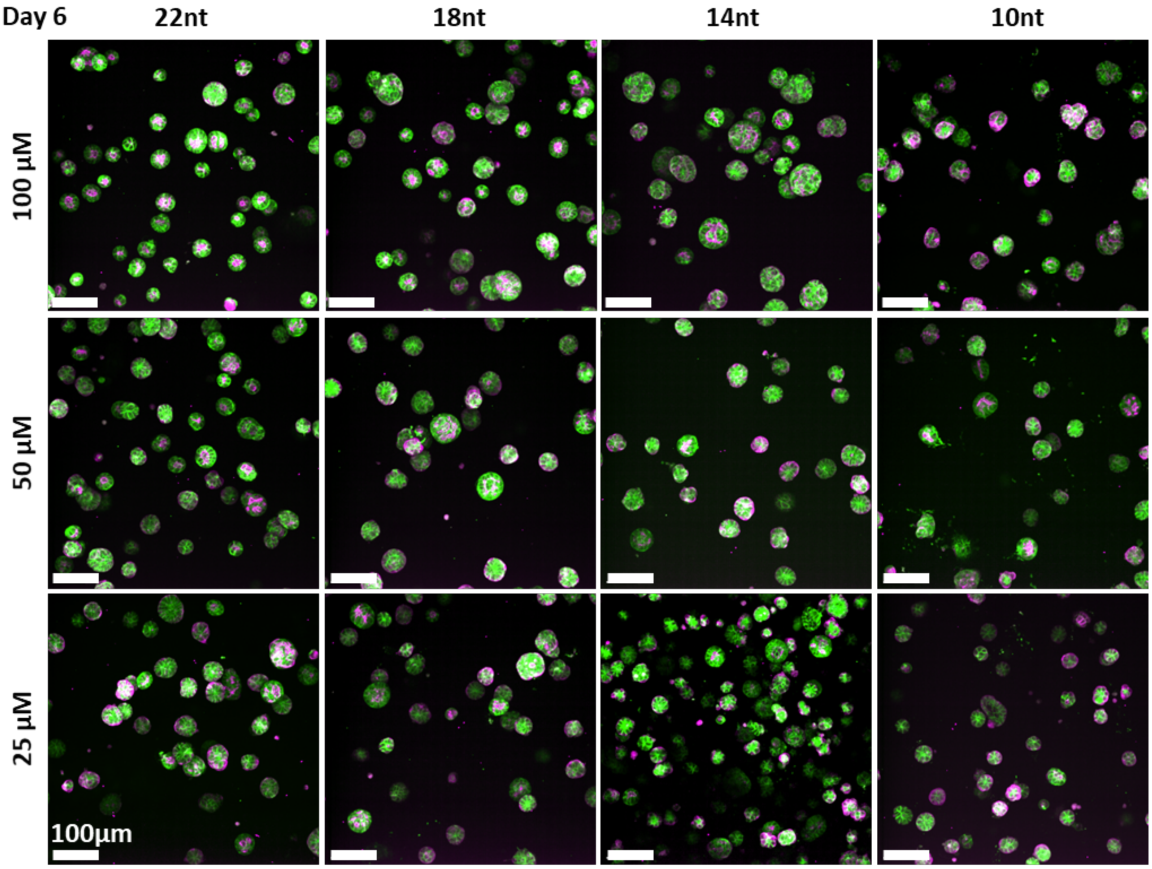

**Figure S7. Confocal images of MDCK cysts in *DyNAtrix* with different stiffness and stress relaxation (day 6).**

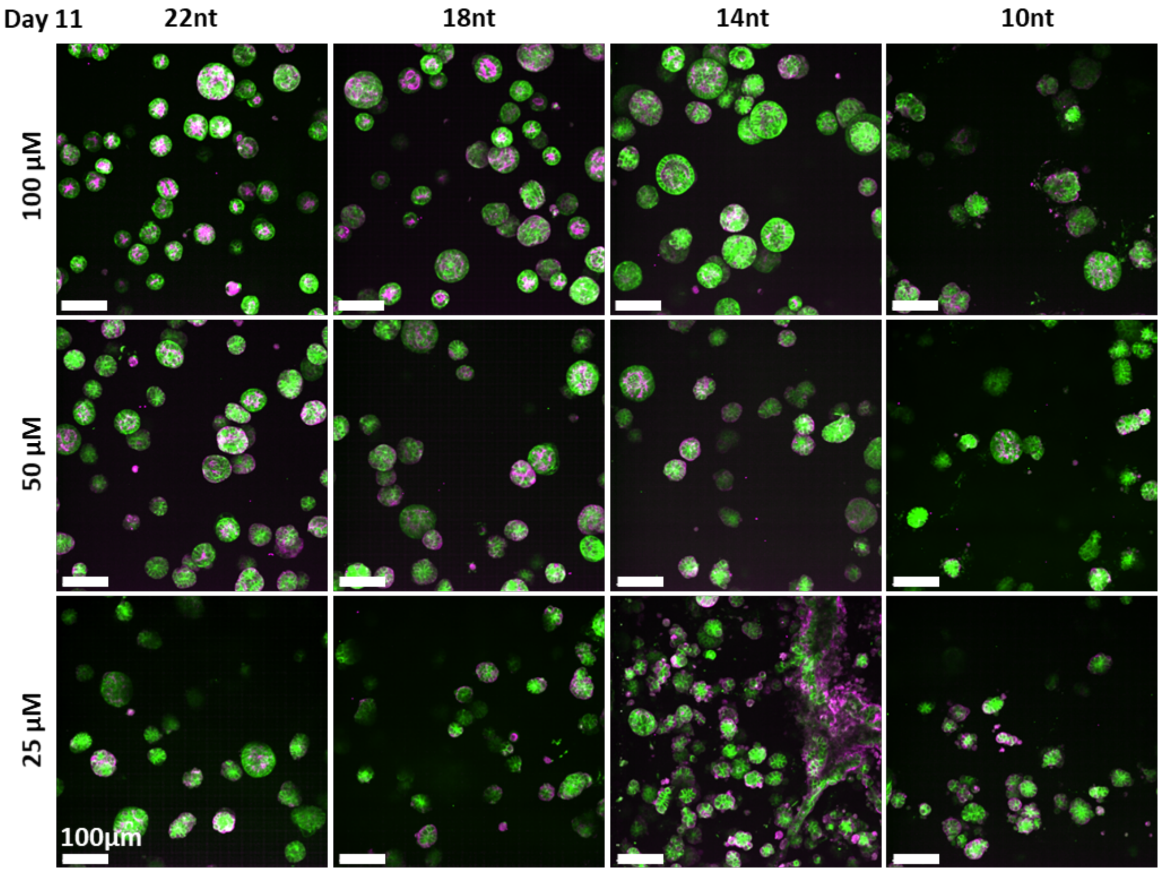


**Figure S8. Confocal images of MDCK cyst cultured in *DyNAtrix* with different stiffness and stress relaxation (day 11).**

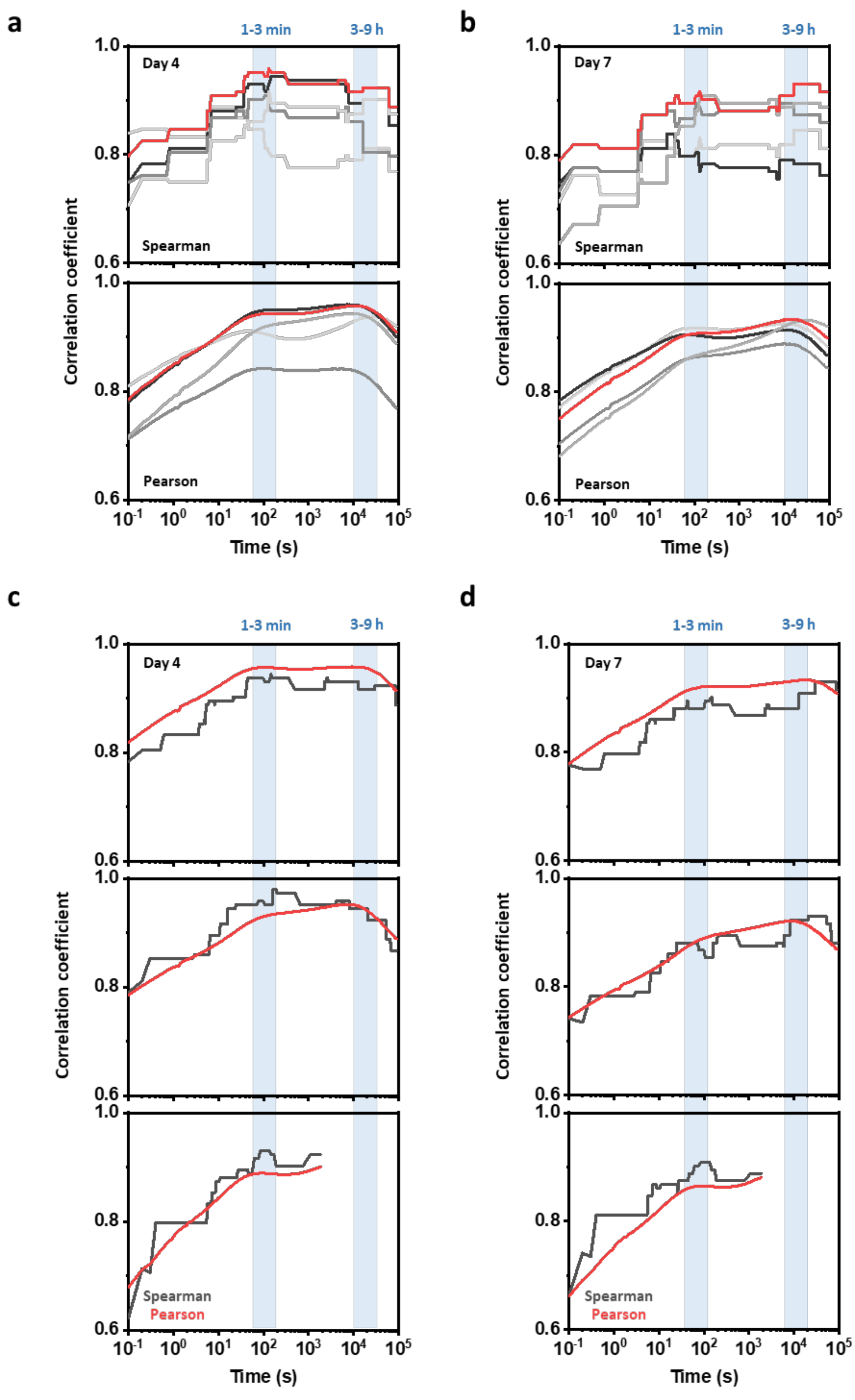


**Figure S9. Correlation analysis of apical-in polarity and shear stress over time. a,** Correlation of individual biological repeats (**day 4**) to the average of the stress-relaxation data. **b,** Correlation of individual biological repeats (**day 7**) to the average of the stress relaxation data. (**a,b:** 4 repeats are shown in black and gray; the correlation of the average of the biological repeats with the stress-relaxation data is shown in red). **c,** Correlation of individual stress-relaxation repeats with average experimental cyst polarity (**day 4**)). **d,** Correlation of individual stress-relaxation repeats with average experimental cyst polarity (**day 7**).

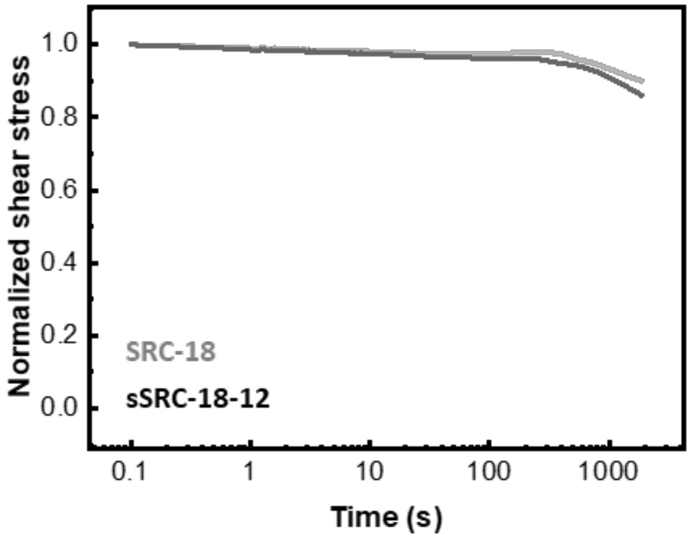


**Figure S10. Normalized stress–relaxation curve of DyNAtrix crosslinked with the SRC-18 or sSRC-18-12.**

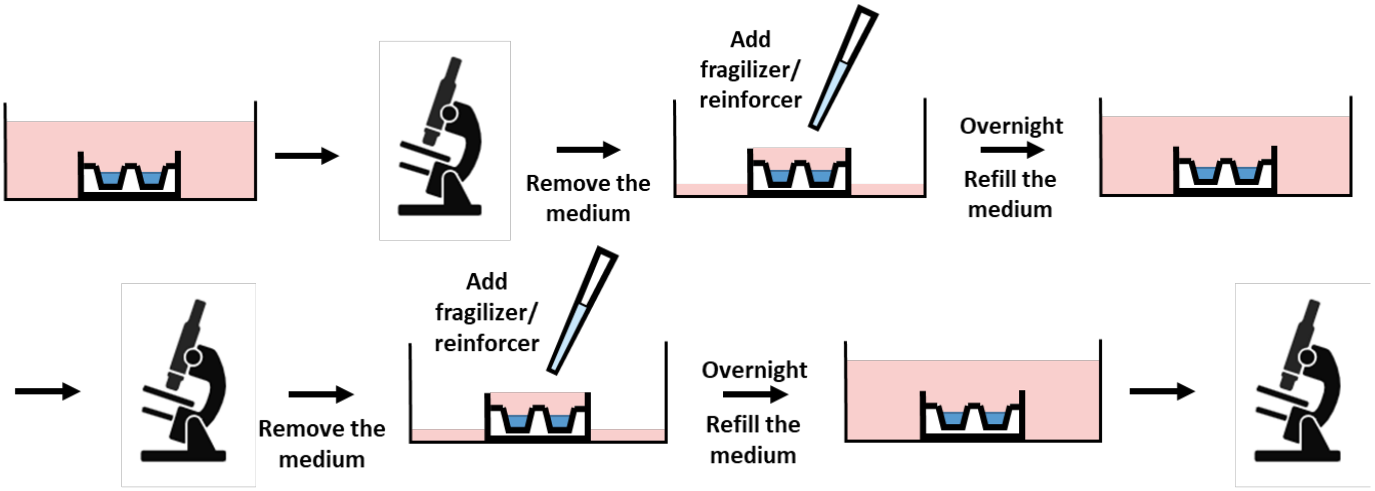


**Figure S11. Scheme of in situ stress-relaxation switching during cell culture.** *DyNAtrix* was placed into micro-well inserts (*ibidi GmbH*). During the switching, most of the medium was removed. Only ~200µL of it was kept in the insert to maximize the concentration of fragilizer and reinforcer. After overnight incubation (16 hours), the medium was refilled.

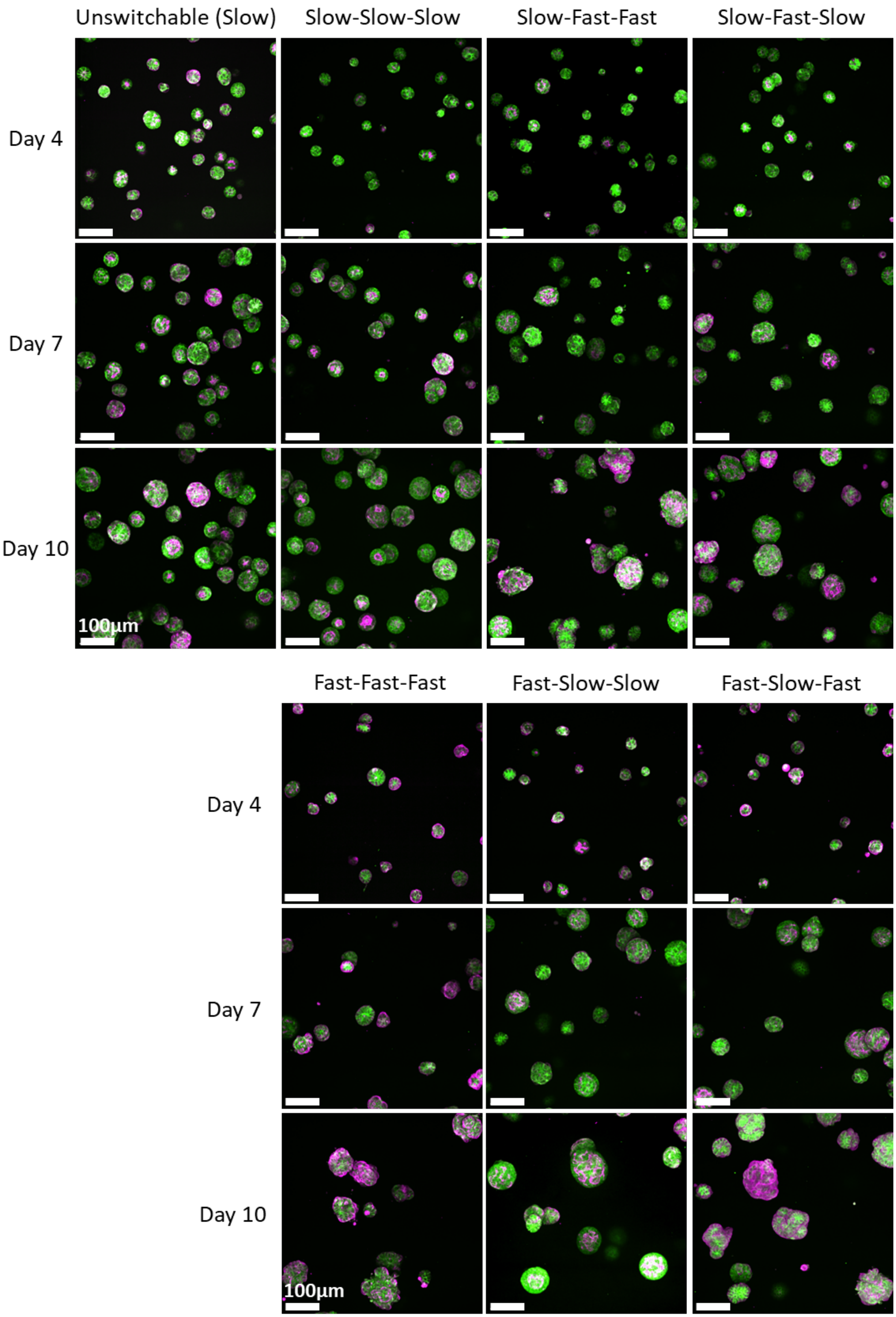


**Figure S12. Confocal images of MDCK cysts that underwent stress-relaxation switching, imaged on days 4, 7, and 10.**

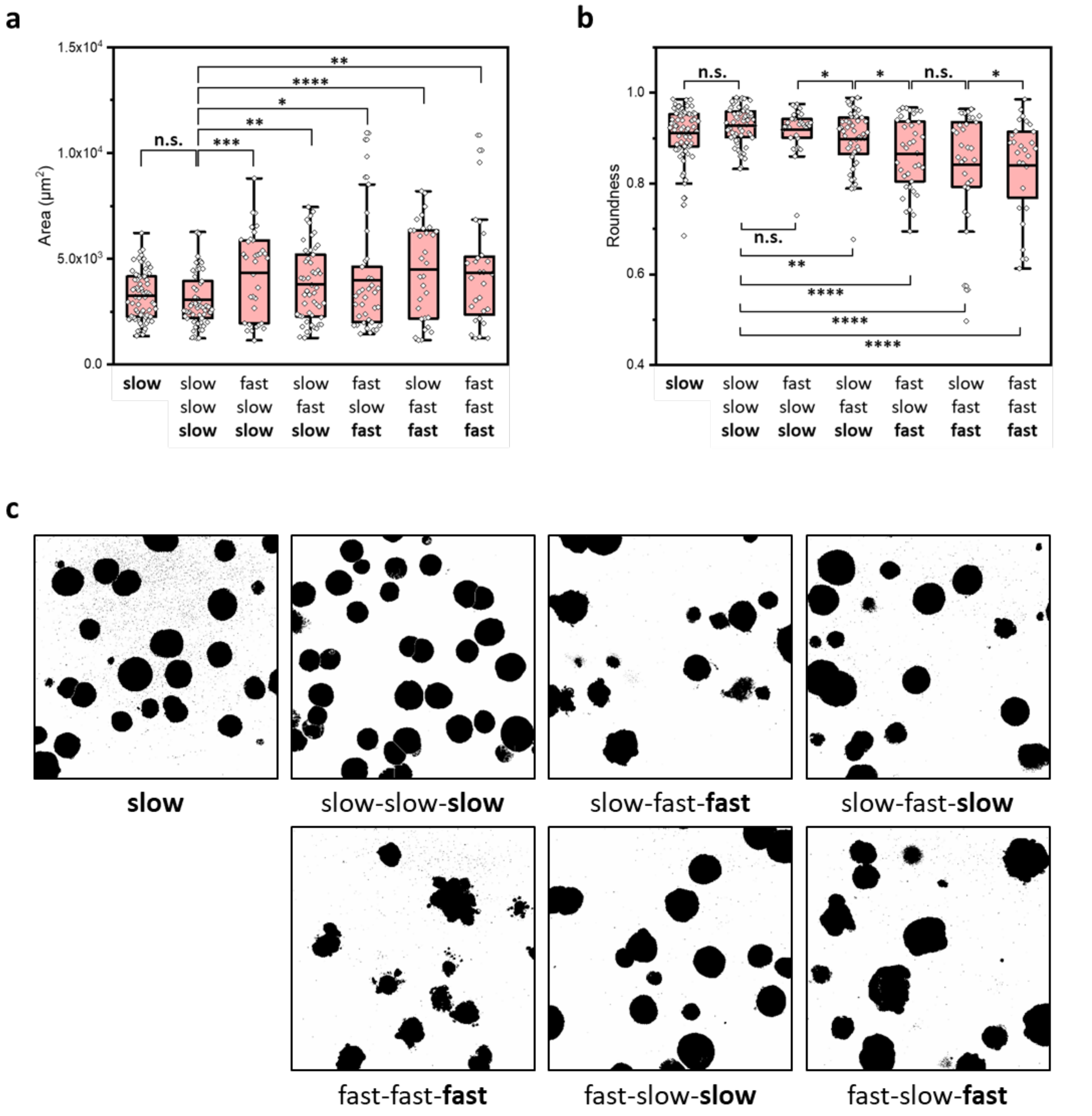


**Figure S13. Analysis of cyst (day 10) size and roundness. a,** Size of cysts in each condition. Size was defined as the projection area of cysts in the confocal images (see panel c). **b,** Roundness of cysts in each condition. **c,** Representative images (maximum intensity projection) of cysts in each condition. Both size and roundness were analyzed in ImageJ. Cysts at the edge of an image and overlapping cysts were excluded from the analysis. Statistical analysis was performed using two-sample t-test (assuming equal variances). n.s., not significant; *, $P < 0.05$; **, $P < 0.01$; ***, $P < 0.001$; ****, $P < 0.0001$.

### Supplementary Tables

**Table S1. List of DNA sequences used in this study.** Red: anchor strand sequence or adaptor site complementary to the anchor strand; blue: crosslinker overlap domain; magenta: ambiguous N base (A,T,C, or G); ochre yellow: binding domain for switchable crosslinker. 5Acryd: Acrydite.

| # | Anchor strand | Length(nt) |
|---|---|---|
| 1 | /5Acryd/GACGGCTCATAAGGCTCTAATC | 22 |

| # | 22nt SRC and its blocker strands | Length(nt) |
|---|---|---|
| 2A | TGTGTTAGTCANTGTCCCATTAGATTAGAGCCTTATGAGCCGTC | 44 |
| 2B | TAATGGGACANTGACTAACACAGATTAGAGCCTTATGAGCCGTC | 44 |
| 2a | ACATGGGACANAGACTAACATT | 22 |
| 2b | AATGTTAGTCTNTGTCCCATGT | 22 |

| # | 18nt SRC and its blocker strands | Length(nt) |
|---|---|---|
| 3A | TGTTAGTCANTGTCCCATGATTAGAGCCTTATGAGCCGTC | 40 |
| 3B | ATGGGACANTGACTAACAGATTAGAGCCTTATGAGCCGTC | 40 |
| 3a | AAGGGACANTGACTAAGT | 18 |
| 3b | ACTTAGTCANTGTCCCTT | 18 |

| # | 14nt SRC and its blocker strands | Length(nt) |
|---|---|---|
| 4A | TTAGTCANTGTCCCGATTAGAGCCTTATGAGCCGTC | 36 |
| 4B | GGGACANTGACTAAGATTAGAGCCTTATGAGCCGTC | 36 |
| 4a | AAGGACANTGACTAGT | 16 |
| 4b | ACTAGTCANTGTCCTT | 16 |

| # | 10nt SRC and its blocker strands | Length(nt) |
|---|---|---|
| 5A | AGTCANTGTCGATTAGAGCCTTATGAGCCGTC | 32 |
| 5B | GACANTGACTGATTAGAGCCTTATGAGCCGTC | 32 |
| 5a | AAGACANTGACTCT | 14 |
| 5b | AGAGTCANTGTCTT | 14 |

| # | sSRC, Fragilizer and Reinforcer | Length(nt) |
|---|---|---|
| 6a | ATGGGACANTGACTAACAGCTGCCAGGCCAGGGATTAGAGCCTTATGAGCCGTC | 54 |
| 6b | GTTATGTGGACCTGGCCTGGCAGCTGTTAG | 30 |
| 6c | CTAACAGCTGCCAGGCCAGGTCCACATAAC | 30 |

| # | Scrambled sequences of Fragilizer and Reinforcer for control experiments | Length(nt) |
|---|---|---|
| 7a | GGTCCAGGTCTAGGACTGTTGGTGGTTACC | 30 |
| 7b | GGTAACCACCAACAGTCCTAGACCTGGACC | 30 |
